# Disruption of the ARID1A-containing SWI/SNF complex reprograms tumor-associated macrophages and enhances immunotherapy response

**DOI:** 10.64898/2026.09.25.754528

**Authors:** Helen M. McRae, Katherine M. Nguyen, Queralt Vallmajó-Martín, Samuel A. Rivera, Braden T. Stevenson, K Garrett Evensen, Mannix J. Burns, Joshua C. Bell, Hannah C. Rattu Mandias, Brent Y. Chick, Isaac Chang, Emily C. Dykhuizen, Kelly Kersten, Susan M. Kaech, Deepshika Ramanan, Diana C. Hargreaves

**Affiliations:** Molecular and Cellular Biology Laboratory, Salk Institute for Biological Studies, La Jolla, CA 92037, USA; NOMIS Center for Immunobiology and Microbial Pathogenesis, Salk Institute for Biological Studies, La Jolla, CA 92037, USA; Biological Sciences Graduate Program, University of California at San Diego, La Jolla, California 92093, USA; Integrative Genomics and Bioinformatics Core, Salk Institute of Biological Studies, La Jolla, CA 92037, USA; Medicinal Chemistry and Molecular Pharmacology, Purdue University, West Lafayette, IN 47907, USA; Cancer Metabolism and Microenvironment Program, Sanford Burnham Prebys Medical Discovery Institute, La Jolla, CA 92037, USA

## Abstract

Tumor-associated macrophages (TAMs) contribute to tumor immune evasion and therapeutic resistance. However, the epigenetic and transcriptional regulators that control TAM function remain largely unidentified. Here we investigated the role of the SWI/SNF chromatin remodeling complex in TAMs and whether disruption of SWI/SNF function in TAMs could improve immunotherapy. Pharmacologic inhibition of SWI/SNF ATPase activity improved the efficacy of checkpoint blockade immunotherapy, slowing tumor growth and reprogramming transcription broadly in tumor cells, tumor-infiltrating lymphocytes, and TAMs. To define the role of SWI/SNF in TAMs specifically, we genetically deleted the SWI/SNF subunit *Arid1a* in myeloid cells and found this was sufficient to suppress tumor progression and enhance checkpoint blockade response. Epigenomic and single cell analyses indicated that SWI/SNF inhibition and ARID1A deletion in TAMs reduced accessibility at enhancers of genes associated with poor prognosis, such as *Spp1,* and increased accessibility at promoters of interferon-stimulated genes (ISGs). CD86 was elevated on ARID1A-deficient TAMs and the enhanced immunotherapy response required CD86 costimulation and CD8+ T cells. These findings establish ARID1A-dependent chromatin remodeling as a determinant of TAM gene expression programs and show that disruption of myeloid SWI/SNF function improves checkpoint blockade immunotherapy via TAM reprogramming.

## Introduction

Tumor-associated macrophages (TAMs) are among the most abundant cells in many tumors and exhibit phenotypic and functional heterogeneity. They promote tumor growth, angiogenesis and immune-suppression, and reduce the efficacy of immunotherapy in addition to other standard-of-care treatments such as radiotherapy and chemotherapy^1^. Emerging strategies to target macrophages include depletion approaches via blocking monocyte recruitment or critical growth receptors such as CSF1R. Alternatively, strategies to reprogram TAMs aim to boost the anti-tumor functions of macrophages and are under clinical investigation in combination with checkpoint blockade immunotherapies^2–5^. Anti-tumor properties of macrophages include direct killing by nitric oxide^6,7^, phagocytosis^8,9^, the recruitment and activation of lymphocytes via chemokine and cytokine production^10^, and antigen-presentation to elicit lymphocyte activation^11–14^. TAMs exhibiting pro-tumor and anti-tumor properties coexist in mouse and human tumors and subtypes can be distinguished in single cell analyses^15–18^. Notably, response to immunotherapy in breast cancer and melanoma clinical cohorts is associated with macrophage production of interferon stimulated genes (ISGs)^15–17,19,20^, including *CXCL9* and *CXCL10*^10^, while poor prognosis is associated with expression of *SPP1* (osteopontin)^17,18^, with the ratio of *CXCL9:SPP1* correlating with outcomes in head and neck squamous carcinoma, lung and colon tumors^21^.

Many of the current clinical targets for tumor macrophage reprogramming involve targeting cell surface receptors^2,3^, such as inhibition PI3Kγ^22^, agonism of CD40^23^ and blockade of the CD47/SIRPa axis^9^. Additionally, there is active interest in elucidating the transcriptional and epigenetic factors that regulate TAM function in tumors^24^. Class IIa histone deacetylases (HDAC 4,5,7,9), which are thought to act as adaptor proteins rather than canonical histone deacetylases,^25^ were identified as targets for macrophage reprogramming^26^. Loss of myeloid-specific expression of TET2, a factor that mediates DNA demethylation, was shown to reprogram macrophages in pre-clinical melanoma models^27^, and TET2-mutant clonal hematopoiesis is associated with improved checkpoint blockade immunotherapy response in solid tumors^28^. MYC deletion in myeloid cells slows growth of mouse melanoma^29^. An *in vitro* screen for macrophage reprogramming factors identified the transcription factor ZEB2 as driving a pro-tumor gene expression program, and knockdown of *Zeb2* using a CpG-siRNA construct reduced tumor growth^30^. These studies suggest that the pro-tumor function of TAMs is regulated by transcriptional and epigenetic mechanisms that can be inhibited to induce TAM reprogramming, but the full suite of nuclear factors that control TAM function is unknown.

Macrophage chromatin accessibility is highly dependent on the local tissue^31^ and tumor microenvironment^32–35^, suggesting chromatin remodeling is required to integrate specific cues from the tumor microenvironment and drive tumor-specific transcriptional programs. SWI/SNF complexes are ATP-dependent nucleosome remodeling complexes that regulate cell type-specific chromatin accessibility. There are three major SWI/SNF complex configurations, the ARID1A-containing canonical BAF (cBAF), non-canonical BAF (ncBAF), and Polybromo-associated BAF (PBAF). SWI/SNF interacts with PU.1 and is essential for myeloid differentiation^36–42^. ARID1A also plays important roles in tissue-resident macrophage populations such as promoting the bone-resorption capacity of osteoclasts^43,44^ and branching of microglia^45^. In bone marrow-derived macrophages (BMDMs), SWI/SNF complexes are recruited to inflammatory genes following endotoxin stimulation and perturbation of the SWI/SNF complex reduces expression of inflammatory response genes^46–48^. However, it is not known whether SWI/SNF regulates differentiation or function of myeloid cells in response to cues in the tumor microenvironment.

Subunits of the SWI/SNF complex frequently undergo loss-of-function mutations in cancer^49^. Interest in therapeutically targeting the SWI/SNF complex has been promoted by the discovery of synthetic lethal relationships between paralogous subunits of SWI/SNF^50^. Even in the absence of SWI/SNF complex mutations, several cancer types require the SWI/SNF complex for their survival, including AML^36^, uveal melanoma^51^, myeloma^52^, B cell lymphoma^53,54^, prostate cancer^55^ and POU2F3+ small cell lung cancer^54,56,57^. Preclinical studies using SWI/SNF targeted therapies have shown efficacy in immune-deficient mouse models^37,51–55,58^, which are agnostic to the effects of these agents on immune cells. Two recent studies using BRM014 (SWI/SNF ATPase inhibitor)^59^ or AU-15330 (SWI/SNF ATPase degrader)^60^ in immune-competent ovarian cancer models observed increased immune infiltration associated with reduced tumor growth. How SWI/SNF pharmacologic inhibition affects tumor-infiltrating immune cells is still unclear.

Here, we show that SWI/SNF inhibition in combination with anti-PD-L1 blockade is effective at reducing tumor growth concomitant with extensive transcriptional changes in TAMs, CD8+ T cells and tumor cells. Using lineage-specific genetic deletion, we found that ARID1A deletion in myeloid cells was sufficient to decrease tumor growth and improve immunotherapy response. ARID1A deletion induced TAM reprogramming, resulting in changes to chromatin accessibility that impacted the overall balance of tumor-promoting and anti-tumor gene expression in TAMs in tumors. Furthermore, tumor reduction in hosts with myeloid-specific *Arid1a* deletion was dependent on CD8+ T cells and the CD86 costimulatory molecule, upregulated on *Arid1a*KO TAMs. This work suggests efficacy of SWI/SNF inhibition combined with checkpoint blockade immunotherapy and highlights a role for the SWI/SNF complex in epigenetic-driven TAM reprogramming.

## Results

### SWI/SNF inhibition enhances tumor control in combination with anti-PD-L1 immunotherapy

To test whether SWI/SNF inhibition could enhance the response to checkpoint blockade immunotherapy, we treated immune-competent mice bearing subcutaneous MC38 colon adenocarcinoma tumors with a combination of the SWI/SNF ATPase inhibitor FHD-286 and anti-PD-L1 starting at day 12 post-tumor injection when control tumors are partially responsive to anti-PD-L1. In combination with daily oral gavage of FHD-286^51^ at 1.5 mg/kg, we observed a significant decrease in tumor growth compared to mice receiving vehicle and anti-PD-L1 (Fig. 1A-C, Supplementary Fig. 1A). A similar reduction in tumor growth was observed with an alternative SWI/SNF ATPase inhibitor, BRM014^61^ (Supplementary Fig. 2A-D). The viability of MC38 cells *in vitro* was not highly sensitive to SWI/SNF inhibitor treatment (Supplementary Fig. 1B), prompting us to examine the contribution of the tumor microenvironment to the effect observed *in vivo.* We performed flow cytometry on the tumor immune infiltrate (Supplementary Fig. 1C) and identified a significant increase in the absolute number of CD8 T cells per mg tumor in FHD-286 versus vehicle treated samples (Fig 1D), but not in other lymphocyte populations (B cells, CD4+ conventional, regulatory T cells and NK cells), myeloid populations (monocytes, neutrophils, and macrophages) (Fig. 1D), nor overall immune infiltrate (Supplementary Fig. 1D). cDC1 dendritic cells were decreased, while cDC2 were unchanged (Figure 1D). Similar effects on the tumor immune infiltrate were observed with BRM014 + anti-PD-L1 compared to vehicle + anti-PD-L1 treatment (Supplementary Fig. 2).

**Fig. 1:**
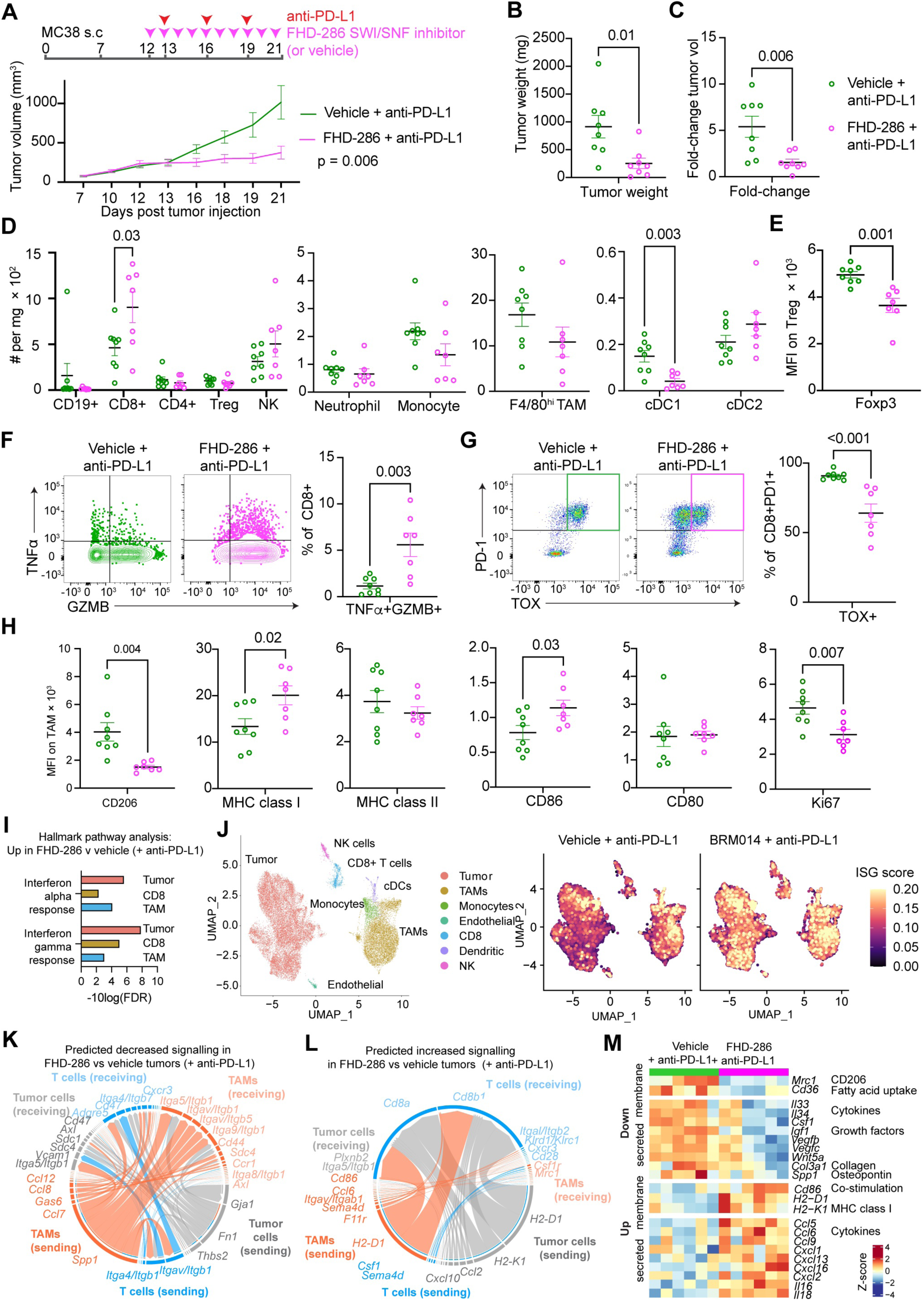
SWI/SNF complex inhibition increases immune activation and alters transcription in the tumor microenvironment in combination with anti-PD-L1 immunotherapy. A. Experimental design showing treatment timepoints and growth (volume) of subcutaneously implanted MC38 tumors into immunocompetent mice treated with FHD-286 and anti-PD-L1 as indicated compared to the those treated with vehicle + anti-PD-L1 (1.5 mg/kg FHD-286 delivered by oral gavage and 200 µg anti-PD-L1 intraperitoneal per dose). B. Tumor weight at day 21 of isolated MC38 tumors of indicated treatment. C. Fold change in tumor volume from first treatment (day 21/day 12). D. Immune infiltrate quantification as assessed by flow cytometry. E. Quantification of Foxp3 protein expression gated on Treg cells (CD45+CD4+FOXP3+). F. Representative staining and quantification of cytokine expression, gated on CD8+ T cells following 4hr PMA + ionomycin stimulation of a single-cell suspension dissociated from MC38 tumors. G. Representative staining and quantification of the proportion of CD8+PD1+ T cells that are expressing high levels of TOX protein. H. Quantification of cell surface markers on TAMs (CD45+CD11b+Ly6C^neg^F4/80+). I. Selected upregulated Hallmark pathways from sorted cells of indicated cell types comparing FHD-286 versus vehicle treatment in the context of anti-PD-L1 treatment (full results in Supplementary Table 2). J. Single-cell multiome analysis of the tumor microenvironment of MC38 tumors showing cell-type annotations and expression of ISGs in the indicated conditions. K. CellChat analysis showing predicted downregulated ligand-receptor interactions between tumor cells, CD8+ T cells and TAM in FHD-286 versus vehicle treatment in the context of anti-PD-L1, highlighting selected interactions. L. CellChat analysis showing predicted upregulated ligand-receptor interactions between tumor cells, CD8+ T cells and TAM in FHD-286 versus vehicle treatment in the context of anti-PD-L1. highlighting selected interactions. M. Heatmap showing row z-scores of selected genes encoding ligand and receptor proteins in TAMs from FHD-286 versus vehicle treated tumors. Data in A analyzed by a linear mixed model using the TumGrowth package with breakpoint at day 12 (treatment initiation) and subjected to a type II ANOVA and pairwise comparisons between conditions. Data in B-H each analyzed by a two-tailed t-test between the groups, with shared legend in panel C. MFI = median fluorescence intensity. Data presented as mean ± SEM, with points indicative of biological replicates.

Corroborating our results from a CRISPR screen for FOXP3 regulators^62^, we found that FOXP3 protein was downregulated in Tregs from FHD-286 + anti-PD-L1 tumors (Fig. 1E, Supplementary Fig. 1E). In addition to increased CD8+ T cell infiltration, we observed an increase in polyfunctional TNFα+GZMB+ CD8+ T cells (Fig. 1F), indicating heightened CD8+ T cell effector function. Strikingly, the proportion of CD8+PD1+ T cells expressing the exhaustion-related transcription factor TOX was reduced by 30% in tumors treated with the combination of FHD-286 + anti-PD-L1 (Fig. 1G), and there was a reduction in the proportion of SLAMF6+TIM3^neg^ “progenitor exhausted” and TIM3+TOX+ “terminally exhausted” cells (Supplementary Fig. 1F-I), consistent with prior reports demonstrating reduced exhaustion in *Arid1a*-deficient intratumoral CD8+ T cells^63^.

In addition to effects on T cells, FHD-286 + anti-PD-L1 had profound effects on cell surface protein expression associated with TAM function, despite having a limited effect on TAM infiltration. We observed a reduction in CD206 (mannose receptor), a marker correlated with increased tumor burden in mice^64^ and an increase in cell surface expression of the antigen-presenting molecule, MHC class I, but not MHC class II, and an increase in the costimulatory protein CD86, but not its paralog CD80 (Fig. 1H, Supplementary Fig. 1J). Intracellular Ki-67 expression was decreased indicating a reduction in TAM proliferation (Fig. 1H, Supplementary Fig. 1J). Thus, inhibition of the ATPase subunits of the SWI/SNF complex causes significant changes to the tumor immune infiltrate composition and activation states, including changes in lymphocytes and TAMs.

## SWI/SNF inhibition transcriptionally alters the tumor microenvironment

To examine the effects of SWI/SNF inhibition on the tumor microenvironment at large, we performed RNA-sequencing (RNA-seq) on whole tumors. We identified 2807 downregulated and 2641 upregulated genes in tumors treated with FHD-286 + anti-PD-L1 versus vehicle + anti-PD-L1 (Supplementary Fig. 3A, Supplementary Table 1). In line with our flow cytometry data, we observed increased gene expression of *Cd8a*, *Gzmb*, and *B2m* (encoding a subunit of the MHC class I complex), while *Foxp3* and *Mrc1* (encoding CD206) were reduced (Supplementary Fig. 3A).

To identify gene expression changes in individual cell types, we performed RNA-seq analysis on sorted tumor cells, CD8+ T cells and TAMs from FHD-286 or vehicle + anti-PD-L1 treated tumors (Supplementary Fig. 3B, Supplementary Table 1). Gene set enrichment analysis revealed that the Interferon Alpha and Interferon Gamma Response pathways were enriched among upregulated genes in tumor cells, TAMs, and CD8+ T cells (Fig. 1I, Supplementary Table 2). This was corroborated by single-cell multiome analysis of the tumor microenvironment following treatment with BRM014 + PD-L1 which indicated widespread upregulation of ISGs (Fig. 1J). Analysis of Interferon Stimulated Genes (ISGs) revealed common and unique upregulated genes in tumor cells, CD8+ T cells and TAMs isolated from FHD-286 treated tumors (Supplementary Fig. 3C). This is consistent with upregulation of ISGs in tumor cells treated with SWI/SNF ATPase inhibitors/degraders *in vitro*, which we and others previously reported^59,60,65,66^. Further, there was a significant positive correlation between gene expression changes in CD8+ T cells from FHD-286 treated tumors and published datasets from *Tox* knockout CD8+ T cells^67^ (Supplementary Fig. 3D) and *Arid1a* knockout intratumoral CD8+ T cells^63^ (Supplementary Fig. 3E), indicating that SWI/SNF inhibition induces similar changes in T cell effector and exhaustion signatures as *Tox*^67^ and *Arid1a*^63^ genetic deletion in CD8+ T cells.

We expected that FHD-286-induced anti-tumor immunity was dependent on interactions between cell types and thus performed CellChat^68,69^ analysis of tumor, CD8+ and TAM populations. The top predicted decreased pathway between FHD-286 and vehicle-treated tumors was the loss of interactions between SPP1 on TAMs and integrins on TAMs, CD8 T cells, and tumor cells (Fig. 1K). SPP1 produced by TAMs is one of the top associations with poor prognosis in human solid tumors^21^. Predicted upregulated interactions in FHD-286 treated tumors were between MHC class I on TAMs and tumor cells with CD8 T cells (Fig. 1L). Other upregulated pathways include the costimulatory CD86 from TAMs interacting with CD8+ T cells (Fig. 1L). These changes were underscored by changes in gene expression in TAMs (Fig. 1M).

These data indicate that SWI/SNF inhibition leads to broad transcriptional changes in the tumor microenvironment, including effects that phenocopy effects of SWI/SNF inhibition in tumor cells^65^ and *Arid1a* deletion in tumor-infiltrating CD8+ T cells^63^ as well as previously undescribed changes in TAMs that comprise gain of expression of ISGs (MHC class I, *Cd86* and chemokines), and reduction in pro-tumor genes such as *Spp1*.

## Deletion of *Arid1a* in myeloid cells leads to reduced tumor growth and heightened immunotherapy response

The effect of systemic SWI/SNF inhibition on TAM cell surface phenotypes and gene expression suggested an important role for the SWI/SNF complex in the regulation of TAM function. To investigate the cell-intrinsic effects of disrupting the cBAF complex in the myeloid lineage, we used an *Arid1a* floxed mouse model^70^ combined with myeloid-specific Cre recombinase drivers^71–73^. We profiled tissue resident macrophages in non-tumor bearing mice and confirmed that ARID1A is highly expressed and efficiently deleted with *LysM^Cre^* (Supplementary Fig. 4A). There were no changes in the proportion or number of tissue resident macrophages in the lung, colon, liver or spleen, indicating that deletion of ARID1A in macrophages is compatible with macrophage survival (Supplementary Fig. 4B-C). As expected, we observed significant deletion of ARID1A in peripheral blood neutrophils with *LysM^Cre^*(Supplementary Fig. 4D). Reporter studies show *LysM^Cre^* expression in peripheral Ly6C^hi^ monocytes^74^; however, we observed loss of ARID1A protein only in the longer-lived Ly6C^neg^ patrolling monocytes (Supplementary Fig. 4D). We observed no changes in the proportion or number of neutrophil or monocyte populations in the peripheral blood (Supplementary Fig. 4E-F).

To test whether *Arid1a* deletion in the myeloid compartment was sufficient to affect tumor growth, we injected MC38 tumors into *Arid1a^f/f^;LysM^Cre^* or control mice and monitored tumors without treatment or following injection of anti-PD-L1 at day 13 (Fig. 2A). As expected, controls were partially responsive to anti-PD-L1 treatment (Fig. 2A). Strikingly, the effect of myeloid-specific ARID1A deletion was similar to the effect of anti-PD-L1 alone, and addition of anti-PD-L1 immunotherapy further increased tumor control in *Arid1a^f/f^;LysM^Cre^* mice (Fig. 2A-B). ARID1A was abundantly expressed in TAMs and efficiently deleted with *LysM^Cre^* (Supplementary Fig. 5A). We extended our analysis to an alternative myeloid Cre driver, *Itgax^Cre^,* which deletes in macrophages and dendritic cells, but not neutrophils^74^ (Supplementary Fig. 5B). We found that tumors grew smaller in untreated *Arid1a^f/f^;Itgax^Cre^*mice compared to untreated control mice (Supplementary Fig. 6A-B). To test whether constitutive ARID1A deletion was required for the observed effect on tumor growth, we deleted ARID1A in MC38 tumor-bearing adults using a tamoxifen-inducible *LysM^CreERT2^* model. *Arid1a^f/f^;LysM^CreERT2^*mice treated with tamoxifen displayed similar tumor reduction (Supplementary Fig. 6C-D), indicating acute deletion of ARID1A after tumors are established is sufficient to slow tumor growth. As expected, myeloid genetic deletion was not as effective as pharmacologic SWI/SNF inhibition at slowing tumor growth in MC38 tumors given that SWI/SNF inhibition affects tumor cells and CD8+ T cells (Fig. 1I, J), among other possibly contributing cell types. In addition, SWI/SNF inhibition will disrupt all SWI/SNF complex types present in all cells, including the remaining PBAF, ncBAF and ARID1B-containing cBAF complex present in ARID1A-deficient myeloid cells.

**Fig. 2:**
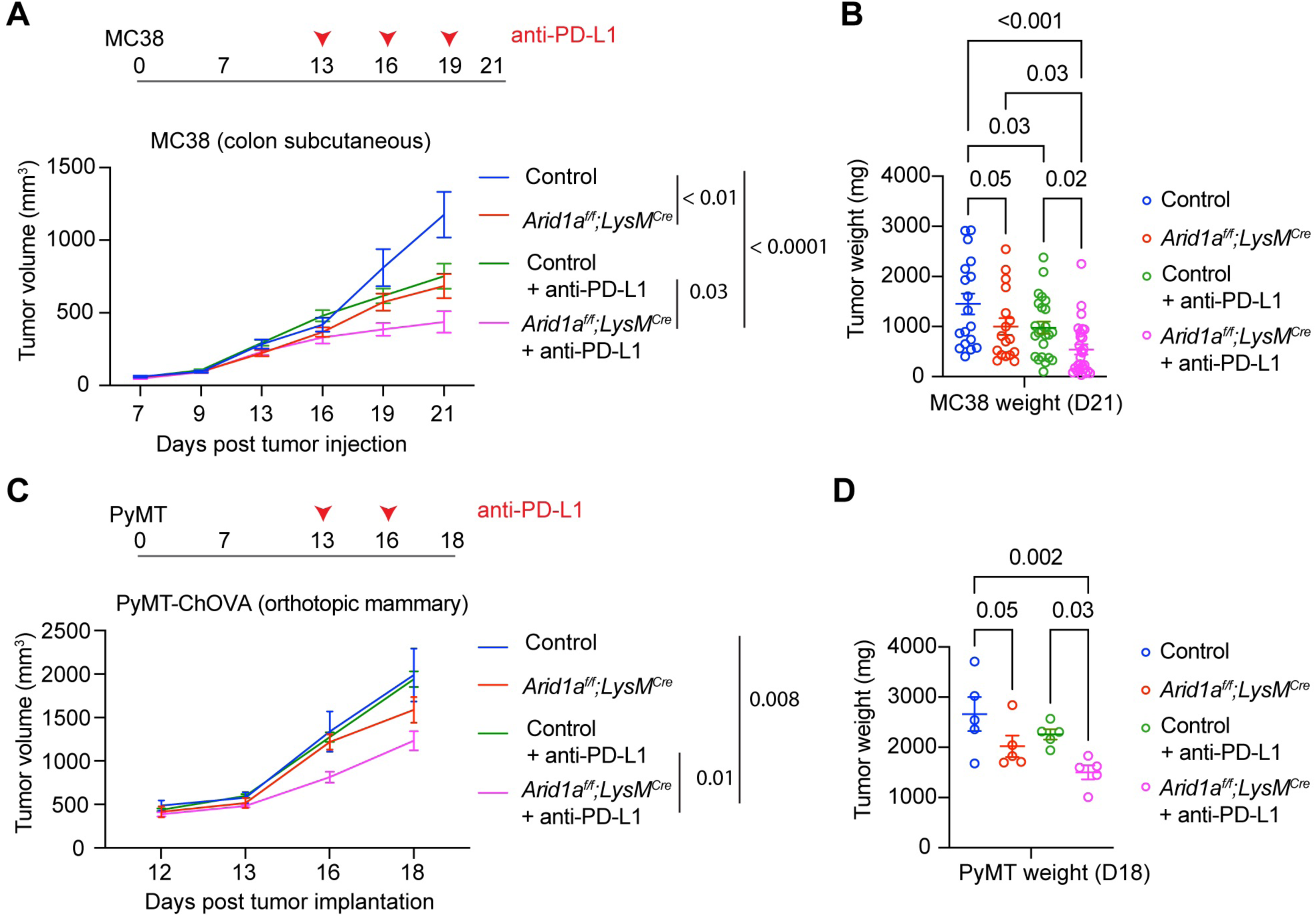
Genetic disruption of the ARID1A-containing SWI/SNF complex in the myeloid lineage leads to slower tumor growth and enhanced immunotherapy response. A. Tumor volume of MC38 subcutaneously implanted tumors into mice lacking myeloid-specific ARID1A (*Arid1a^f/f^;LysM^Cre^*) versus controls, either untreated (isotype-only) or anti-PD-L1 treated. B. Tumor weight at day 21 of MC38 subcutaneous tumors. C. Tumor volume of PyMT-ChOva cell line orthotopically implanted into the mammary fat pad of the indicated genotypes, either untreated or treated with anti-PD-L1. D. Tumor weight at day 21 or PyMT-ChOva orthotopic tumors. Data in A analyzed by a linear mixed model using the TumGrowth package with breakpoint at day 13 (treatment initiation) and subjected to a type II ANOVA for comparisons between conditions. Data in B showing results of multiple comparison testing following ANOVA. Each data point is a biological replicate and error bars are SEM.

Next, we sought to determine the applicability of our findings in orthotopic tumors that are insensitive to checkpoint blockade immunotherapy. We surgically implanted PyMT-ChOVA^75^ into the mammary fat pad of *Arid1a^f/f^;LysM^Cre^* and control mice. Consistent with previous data, the PyMT-ChOVA line was resistant to blockade of the PD-1:PD-L1 axis in controls^76^. However, we observed reduced tumor growth in *Arid1a^f/f^;LysM^Cre^* hosts and increased tumor control in combination with anti-PD-L1 treatment (Fig. 2C-D). Similar findings were observed with subcutaneous Lewis Lung Carcinoma (LLC) tumors in untreated *Arid1a^f/f^;LysM^Cre^*mice (Supplementary Fig. 6E-F). Together, these data show that cell-intrinsic deletion of ARID1A in myeloid cells is sufficient to reduce tumor growth and enhance checkpoint blockade response in immunogenic and immunotherapy insensitive tumor models.

## Myeloid-specific *Arid1a* deletion promotes TAM reprogramming

To profile the cellular changes in TAMs lacking ARID1A, we performed immunophenotyping by flow cytometry of untreated or anti-PD-L1-treated MC38 tumors from control and *Arid1a^f/f^;LysM^Cre^*mice (Supplementary Fig. 7A). There was no change in the absolute number of infiltrating immune cells or *Arid1a*KO (*Arid1a^f/f^;LysM^Cre^*) TAMs per mg in MC38 tumors and only modest changes in other myeloid populations (Supplementary Fig. 7B-C). However, we identified changes in phenotypic markers on *Arid1a*KO TAMs consistent with reprogrammed function. *Arid1a*KO TAMs displayed increased PD-L1 expression (Fig. 3A, Supplementary Fig. 7D) and decreased CD206 (Fig. 3B, Supplementary Fig. 7E). MHC class I, MHC class II, and SLAMF7 were increased on *Arid1a*KO TAMs from anti-PD-L1 treated *Arid1a^f/f^;LysM^Cre^* mice, while CD86 was increased on *Arid1a*KO TAMs from both untreated and anti-PD-L1 treated groups (Fig. 3B, Supplementary Fig. 7E). *Arid1a*KO TAMs had increased expression of lineage markers F4/80 and CD11b, but no change in CD64, CD11c, or CD80 (Supplementary Fig. 7F-G). Thus, macrophage lineage identity is largely preserved in the context of ARID1A deletion, with changes in the levels of functional cell surface proteins.

**Fig. 3:**
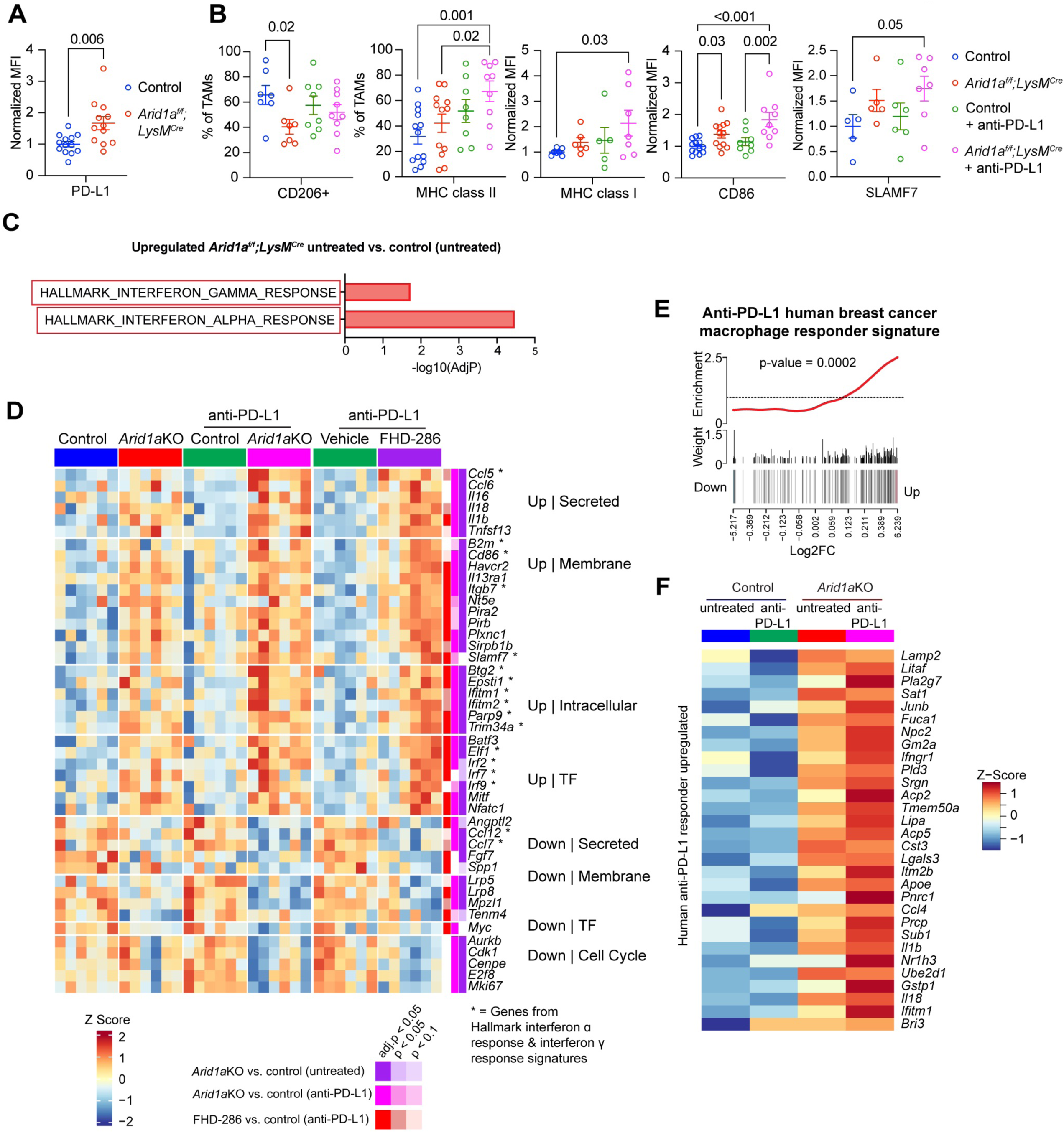
Loss of ARID1A reprograms TAMs. A. Quantification of PD-L1 expression in untreated tumors of the indicated genotypes (representative histogram in Supplementary Figure 7D) B. Quantification of the indicated cell surface receptors on TAMs of indicated genotypes/treatment from MC38 tumors (quantification as percentage where expression is bimodal and as MFI where expression follows a normal distribution, see Supplementary Figure 7E). MFI = median fluorescence intensity, normalized to the control group in each experiment where data from more than one experiment was combined. Data presented as mean ± SEM, with points indicative of biological replicates. C. Hallmark interferon pathways significantly increased in expression in *Arid1a^f/f^;LysM^Cre^*versus control TAM comparisons from either untreated (isotype-only) treated tumors (full results for all conditions and barcode plots in Supplementary Figure 8E). D. Heatmap showing selected genes across biological replicates encoding ISGs, secreted proteins, membrane receptors, transcription factors, and cell cycle genes with significance level in the 3 datasets (*Arid1a^f/f^;LysM^Cre^*versus controls untreated, *Arid1a^f/f^;LysM^Cre^*versus controls anti-PD-L1, and FHD-286 versus vehicle anti-PD-L1) indicated by color on the right. E. Barcode plot showing positive correlation with Zhang et al.^17^ macrophage anti-PD-L1 responder signature from human breast cancer patients. F. Heatmap of commonly upregulated genes in Zhang et al.^17^ macrophage responder signature and *Arid1a^f/f^;LysM^Cre^* versus control TAMs, showing the average z-score from 6 biological replicates for each condition. Data in B and D each analyzed by one-way ANOVA. Error bars are SEM.

To examine how loss of ARID1A affected the overall transcriptional landscape of TAMs, we performed RNA-seq. We identified 677 significantly downregulated and 691 upregulated genes between *Arid1a*KO versus control TAMs from isotype-treated MC38 tumors (Supplementary Fig. 8A, Supplementary Table 3), and 511 downregulated and 618 upregulated between genotypes from anti-PD-L1 treated tumors (Supplementary Fig. 8B, Supplementary Table 3). Gene expression changes between *Arid1a*KO and control TAMs were similar, regardless of anti-PD-L1 treatment (Supplementary Fig. 8C, r = 0.86). Furthermore, gene expression changes in *Arid1a*KO TAMs and TAMs from FHD-286 treated tumors were positively correlated (Supplementary Fig. 8D, r = 0.62). Similar to our results with FHD-286 (Fig. 1J), the Interferon Alpha and Gamma Response were the most significantly enriched pathways among upregulated genes in *Arid1a*KO TAMs (Fig. 3C, Supplementary Fig. 8E), while Myc targets were downregulated (Supplementary Fig. 8E). Immune dictionary analysis^77^ for macrophage response to cytokines also identified overrepresentation of the Type I Interferon response pathways in *Arid1a*KO TAMs and TAMs from FHD-286 treated tumors (Supplementary Fig. 8F). These results were surprising in light of our previous work demonstrating a requirement for SWI/SNF remodeling in the upregulation of inflammatory response genes^46^, including ISGs, in response to endotoxin stimulation of BMDMs, indicating the effects of long-term SWI/SNF perturbation of myeloid cells in the tumor microenvironment are distinct from acute effects on *in vitro* stimulation.

Upregulated ISGs in *Arid1a*KO TAMs included genes encoding secreted factors *(Ccl5, Il1a, Il1b, Il16* and *Il18),* cell-surface proteins (*Cd86, Slamf7),* as well as transcription factors (*Irf2, Irf7, Elf2, Batf3*, *Mitf* and *Nfatc1)*, several of which were also differentially expressed with FHD-286 (Fig. 3D). Downregulated genes include growth factors such as *Fgf7* and *Spp1* (Fig. 3D). We also observed decreased expression of *Myc* and several cell cycle genes in response to genetic loss of ARID1A or FHD-286 inhibition (Fig. 3D).

Comparing the effect of ARID1A deletion in TAMs with published TAM-reprogramming gene signatures, we identified a positive correlation of *Arid1aKO* TAMs from anti-PD-L1 treated tumors with PI3Kγ-deficient (*Pik3cg*KO) TAMs^22^ (LLC tumors), TAMs treated for 16h with a CD40-agonist + anti-CSF1R (MC38 tumors)^23^ and *Zeb2*-deficient TAMs (from siRNA plus CpG treated MC38 tumors)^30^ (Supplementary Fig. 8G). On the other hand, there was no correlation with the signature of *Irf8-KO* TAMs from PyMT tumors which display impaired antigen-presentation (r = 0.05, Supplementary Fig. 8G). Genes upregulated in more than one dataset that were also upregulated in *Arid1a*KO TAMs or TAMs from FHD-286 treated tumors included *Ccl5*, *Plac8* and *Ifitm1,* while common downregulation was observed for genes including *Serpinf1, Ccr2*, *Fcrls,* and *Cd93* (Supplementary Fig. 8H).

To investigate whether the gene signature caused by ARID1A loss in TAMs was correlated with outcomes in human cancers, we compared our dataset to genes upregulated in a human TAM “responder” signature in triple-negative breast cancer^17^. We identified a positive correlation (Fig. 3E, p-value = 0.0002), with common upregulation of acid phosphatase gene *Acp5* (TRAP), involved in dephosphorylation of SPP1^78^, *Cst3* involved in antigen processing, *Lipa,* involved in breakdown of fatty acids in the lysosome, cytokines IL-18 and IL-1B, and interferon response genes *Ifngr1* and *Ifitm1* (Fig. 3F). This indicates that loss of myeloid-specific *Arid1a* leads to broad transcriptional changes in TAMs that encompass elements of previously described anti-tumor TAM-reprogramming gene signatures and human-responder signatures, including the interferon stimulated gene signature.

## Deletion of ARID1A leads to decreased accessibility at enhancer regions and increased accessibility at promoters in TAMs

To understand if the chromatin remodeling function of the ARID1A-containing cBAF complex underlies the observed transcriptional and phenotypic effects of ARID1A loss in TAMs, we profiled chromatin accessibility using ATAC-seq. Several thousand sites were significantly up- and downregulated in *Arid1a*KO versus control TAMs sorted from untreated or anti-PD-L1 treated MC38 tumors (Supplementary Fig. 9A). Differentially accessible sites in *Arid1a*KO versus control TAMs from isotype and anti-PD-L1 treated tumors were positively correlated (Fig. 4A). We therefore investigated a set of consensus sites sensitive to ARID1A loss, comprising 3639 sites with decreased accessibility and 5328 sites with increased accessibility (Fig. 4A). ARID1A binding (based on ChIP-seq data from BMDMs) was highly enriched at accessible sites, and the majority of increased and decreased accessible sites were bound by ARID1A (Fig. 4B). Although promoters comprise just 20% of all accessible sites, 75% of sites with increased accessibility were located at promoters, with 25% at putative enhancers (Fig. 4C). In contrast, only 2% of sites with decreased accessibility were annotated to promoters while 98% were at putative enhancers (Fig. 4C). Aligning with the fact that promoters are generally more accessible than enhancers, sites with increased accessibility had higher overall accessibility than sites with decreased accessibility in *Arid1a*KO TAMs (Fig. 4C). We profiled H3K4me3 (promoters), H3K4me (enhancers), H3K27ac (active regulatory elements), and H3K27me3 (silenced chromatin) in TAMs from anti-PD-L1 treated MC38 tumors (Fig. 4C-D). Based on these histone modifications, 33% of decreased sites were classified as poised enhancers (H3K4me alone), and 21% were defined as active enhancers by co-occurrence of H3K4me and H3K27ac (Fig. 4C-D). Sites with increased accessibility proximal to a TSS were almost all marked by H3K4me3 (98%), with 86% defined as active by H3K27ac (Fig. 4C-D). Promoters with increased accessibility in *Arid1a*KO TAMs included the promoters of several ISGs that were also transcriptionally upregulated in *Arid1a*KO TAMs (Supplementary Fig. 9D). These data indicate that ARID1A loss in TAMs leads to increased accessibility at active promoters with high basal accessibility and decreased accessibility at poised and active enhancers. Similar results were observed in TAMs from MC38 tumors treated with the ATPase inhibitor BRM014 + anti-PD-L1, in our single-cell multiome dataset (Fig. 4C, Supplementary Fig. 9B-C).

**Fig. 4:**
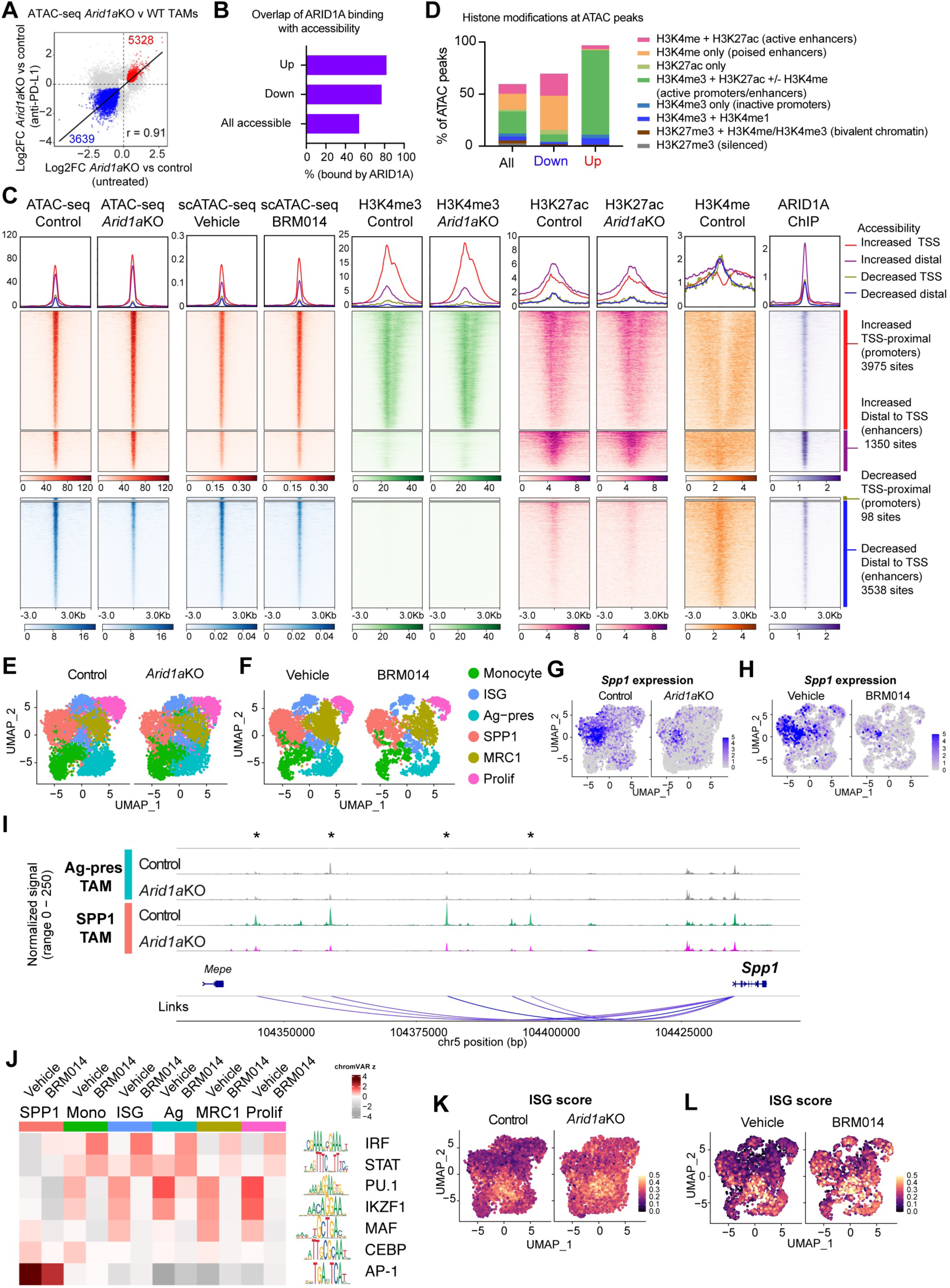
ARID1A loss reprograms chromatin accessibility in TAMs A. Correlation plot showing differential accessibility between peaks in *Arid1a^f/f^;LysM^Cre^* versus control TAMs from untreated or anti-PD-L1 treated conditions, highlighting the consensus set of peaks with decreased accessibility (blue) and increased accessibility (red). Consensus peak set defined as log2FC in the same direction with adjP < 0.05 in at least one condition and p < 0.05 in both conditions. B. Quantification of ARID1A binding at all accessible peaks compared to differentially accessible peaks of each class, based on ARID1A ChIP-seq signal in BMDMs from published dataset^46^. C. Heatmap showing the consensus set of differentially accessible regions between *Arid1a^f/f^;LysM^Cre^*versus control TAMs divided into clusters based on direction of change and distance to nearest TSS element, showing coverage (normalized per cell) in scATAC-seq data of TAMs from anti-PD-L1 + SWI/SNF inhibitor (BRM014) treated and anti-PD-L1 + vehicle-treated MC38 tumors and the levels of indicated histone modifications (from CUT&RUN in TAMs from anti-PD-L1 treated MC38 tumors) and ARID1A binding (from ChIP-seq in BMDM^46^) mapped at those same regions. D. Quantification of histone modification signal at all accessible peaks and differentially accessible peaks. E. UMAP showing TAM substates in MC38 tumors from *Arid1a*KO (*Arid1a^f/f^;LysM^Cre^*) versus control mice (anti-PD-L1 treatment condition). Legend as in (F). F. UMAP showing TAM substates in MC38 tumors from BRM014 versus vehicle treated mice (anti-PD-L1 treatment). G. Expression of *Spp1* transcripts in *Arid1a*KO versus control TAM subsets, showing loss most pronounced in the SPP1 TAM cluster. H. Expression of *Spp1* transcripts in TAMs from BRM014 versus vehicle-treated tumors, showing loss most pronounced in the SPP1 TAM cluster. I. Chromatin accessibility at *Spp*1 enhancers, showing reduced accessibility in *Arid1a*KO versus control genotype SPP1 TAMs (antigen-presenting subset which shows lower accessibility in both groups shown for comparison). J. ChromVAR analysis showing enrichment of motifs across clusters and conditions, showing the top differential motif families between TAM clusters. K. ISG signature score of ISG upregulated in TAMs from *Arid1a*KO versus control tumors, indicating broad increases across TAM subsets. L. ISG signature score of ISG upregulated in TAMs from BRM014 versus vehicle-treated tumors, indicating broad increases across TAM subsets.

## Single-cell analysis reveals chromatin accessibility underlying TAM subtypes and TAM subtype-specific functions of the SWI/SNF complex

SWI/SNF complexes are critical for cell differentiation and loss of SWI/SNF subunits often results in failure to differentiate or loss of cell type-specific gene expression due to the requirement for SWI/SNF-dependent remodeling for lineage TFs^41,42,79,80^. Our bulk analyses point to transcriptional changes upon ARID1A disruption that may reflect changes in overall TAM composition or TAM subtype-specific gene expression, which we sought to unravel using single cell analyses. Several macrophage subtypes have been reported in mouse and human solid tumors including interferon-responsive macrophages, MHC Class II antigen-presenting TAMs, hypoxic/angiogenic/*Spp1* TAMs, *C1q* (complement-producing)/*Mrc1* (CD206) macrophages, and proliferating TAMs^15,16,18,20,21,30,81^, including in MC38 subcutaneous tumors^82,83^. TAM subtypes are correlated with outcomes in human cancers^15–18,20^. In particular, the ratio of *Spp1:Cxcl9* expression, representative of SPP1 and ISG subtypes, was found to predict survival in human dataset^21^.

To assess the effect of ARID1A deletion and SWI/SNF inhibition on TAM heterogeneity, we integrated the monocyte/TAM populations from 3 independent single-cell experiments (comprising data from single-cell RNA and single-nuclei Multiome for simultaneous profiling of gene expression and chromatin accessibility). Clustering subdivided the monocyte/TAM population into 6 cell subtypes: Ly6C^hi^ monocytes (Monocyte), ISG TAMs, MHC class II antigen presenting TAMs (Ag-pres), SPP1 TAMs, MRC1 TAMs and a proliferating cluster (Prolif) (Fig. 4E-F, Supplementary Fig. 10A-B), similar to those previously described in the literature^82^ and displaying overlap with modules identified across human monocyte/macrophages in healthy and tumor tissue^30^ (Supplementary Fig. 10C-D). Trajectory inference using Slingshot pseudotime analysis predicted a pathway with root/terminal nodes at the monocyte and proliferating TAM populations, fitting with a model where TAMs both differentiate from monocytes and are maintained through local proliferation (Supplementary Fig. 10E). Overall, we observed minor changes in TAM composition (Supplementary Fig. 10F). However, when we analyzed subtype-specific accessible sites, we found that sites in the SPP1 and MRC1 clusters were most sensitive to ARID1A loss and BRM014 (Supplementary Fig. 10G), indicating specificity in the role of the cBAF complex across TAM subtypes. Signature genes in the SPP1 and MRC1 subsets were downregulated in BRM014-treated TAMs (Supplementary Fig. 10H). As an example, expression of *Spp1* was reduced in SPP1 cluster (Fig. 4G-H), correlating with reduced accessibility at enhancers in both ARID1A-deleted and BRM014 treated TAMs (Fig. 4I, Supplementary Fig. 10I).

To predict transcription factors regulating TAM subtypes, we first used ChromVar to identify motifs that were most enriched in each TAM cluster. This analysis revealed high enrichment of AP-1 and CEBP family motifs in SPP1 TAMs, MAF-related motifs in MRC1 TAMs, ETS family motifs such as PU.1 (Spi1) in the antigen-presenting and proliferating subclusters of TAMs, and IKZF1 in proliferating TAMs (Fig. 4J, Supplementary Fig. 10J). Accessibility at these subset-enriched motifs was reduced with both BRM014 or ARID1A loss (Fig. 4J, Supplementary Fig. 10J), suggesting that transcription factors previously described to cooperate with the SWI/SNF complex including AP-1^84–86^, CEBP^87,88^, PU.1^36,38–40^ and IKAROS^89^ underlie TAM subtype-specific chromatin accessibility and the dependence on SWI/SNF for subtype-specific gene expression programs. In contrast, IRF and STAT factor motifs were most accessible in monocytes, ISG-TAMs and antigen-presenting TAMs and displayed increased enrichment with SWI/SNF disruption in all TAM subsets (Fig. 4J, Supplementary Fig. 10J). We observed a broad upregulation of ISGs across all clusters, indicating cluster-agnostic ISG derepression in *Arid1a*KO and BRM014 treated TAMs (Fig. 4K-L).

Together, these data demonstrate a role for the SWI/SNF complex in the maintenance of TAM heterogeneity and pro-tumor TAM subtypes. Specifically, ARID1A disruption results in a failure to maintain accessibility and gene expression critical for pro-tumor SPP1 and MRC1 TAMs, while increasing chromatin accessibility and ISG priming in all TAM subtypes. Thus, the ARID1A-containing cBAF complex is a key determinant of TAM phenotypic heterogeneity in tumors through regulation of chromatin accessibility impacting gene expression programs.

### Macrophage-autonomous effects contribute to upregulation of ISGs in *Arid1a*KO macrophages

We previously reported that ARID1A loss in cancer cells and fibroblasts results in ISG upregulation via increased type I interferon production^65^. To determine whether the increase in ISGs was driven by similar macrophage-intrinsic autocrine signaling, we used *in vitro* derived BMDM. We validated that CD86 and PD-L1 were upregulated on *Arid1a*KO BMDM (Fig. 5A). Tamoxifen-inducible ARID1A deletion (*UBC^CreERT2^*) as well as treatment with an ARID1A-inhibitor BRDK98^90^ for 24 hours also increased CD86 and PD-L1 levels (Fig. 5B-C), indicating disruption of ARID1A is sufficient for upregulation. We used an antibody to block the type I interferon receptor (IFNAR) and inhibited JAK1 and JAK2 using ruxolitinib and found the upregulation of CD86/PD-L1 was independent of these factors (Fig. 5D). To test for dependence on all soluble factors, we co-cultured CD45.2+ *Arid1a*KO BMDM with CD45.1+ wildtype BMDM and found that CD86/PD-L1 was exclusively upregulated in the CD45.2+ *Arid1a*KO BMDM, and not on co-cultured wildtype macrophages (Fig. 5E). This indicates cell-intrinsic factors contribute to the upregulation of ISGs in *Arid1a*KO macrophages, distinguishing it from the IFNAR-dependent ISG induction described following loss of ARID1A in cancer cells and fibroblasts^65,91^.

**Fig. 5:**
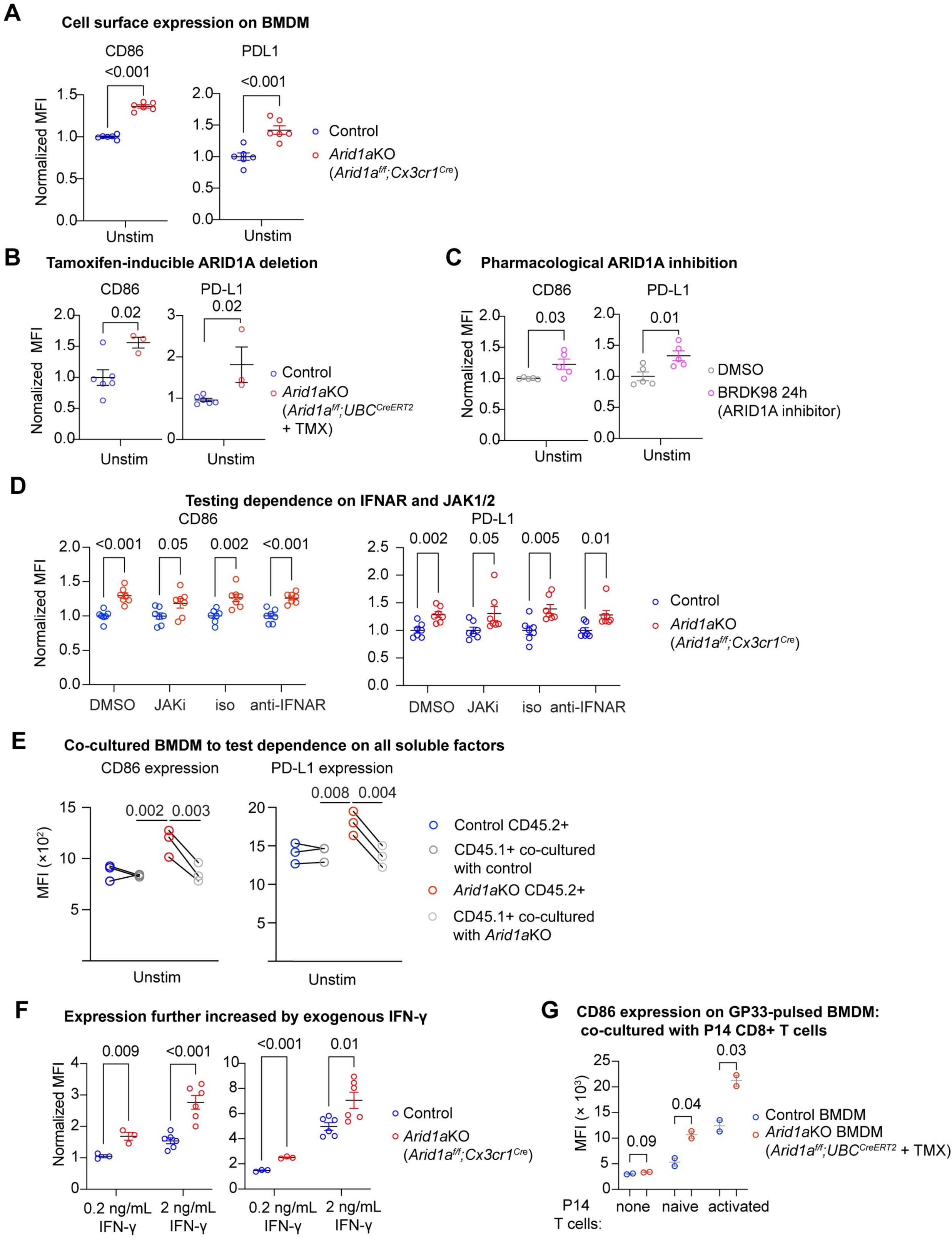
Loss of ARID1A in macrophages leads to cell-intrinsic upregulation of CD86 and PD-L1 that is further inducible by interferon A. Quantification of cell surface CD86 and PD-L1 expression on *Arid1a*KO (*Arid1a^f/f^;Cx3cr1^Cre^*) BMDM relative to control. B. Normalized CD86 and PD-L1 expression following inducible *Arid1a* deletion *in vitro* (tamoxifen-treated *Arid1a^f/f^;UBC^CreERT2^* versus control macrophages). Controls include *Arid1a^f/f^;UBC^CreERT2^* + EtOH, and tamoxifen-treated *Arid1a^+/+^;UBC^CreERT2^*). C. Normalized CD86 and PD-L1 expression following 24h treatment with BRDK98, ARID1A inhibitor. D. CD86 and PD-L1 expression in *Arid1a*KO (*Arid1a^f/f^;Cx3cr1^Cre^*) BMDM relative to control showing upregulation in unstimulated BMDM treated and BMDM treated with JAK1 and JAK2 inhibitor (ruxolitinib) or type I interferon receptor blocking antibody (anti-IFNAR). E. CD86 and PD-L1 expression in co-cultured *Arid1a*KO (*Arid1a^f/f^;Cx3cr1^Cre^*) BMDM (CD45.2+) with wildtype (WT) BMDM (CD45.1+) shown next to with control CD45.1+:CD45.2+ co-culture for comparison. F. Quantification of cell surface CD86 and PD-L1 expression on *Arid1a*KO (*Arid1a^f/f^;Cx3cr1^Cre^*) BMDM relative to control, either unstimulated (unstim) or treated with indicated concentration of IFN-γ. G. CD86 expression following addition of naïve P14 CD8+ T cells or CD3/CD28 pre-stimulated P14 CD8+ T cells to GP33-pulsed BMDM. Data in A-E analyzed by a two-tailed T-test between conditions and data in F analyzed by a paired t-test, grouping those that were co-cultured in the same well. Error bars are SEM and datapoints represent biological replicates.

We wondered if the cell-intrinsic upregulation of ISGs in *Arid1a-*deleted macrophages could be further induced by extrinsic factors present in tumor such as type II interferon from CD8+ T cells. IFN-γ addition further increased CD86 and PD-L1 in a dose-dependent manner (Fig. 5F). Furthermore, co-culture of GP33 peptide-pulsed BMDMs with naïve or preactivated CD8+ T cells expressing the cognate P14 TCR led to hyperinduced CD86 expression in *Arid1a*KO BMDM (Fig. 5G). These data suggest that genetic or pharmacologic inhibition of ARID1A results in basal elevation of CD86 and PD-L1, which is further induced by exogenous factors. Together, these data indicate that ARID1A loss primes macrophages to T cell-derived signals like IFNγ, features that in vivo could engage in a feedforward loop with activated CD8+ T cells to sustain an immunogenic tumor microenvironment.

## Myeloid-specific *Arid1a* deletion increases CD8+ T cell activation

The increase in antigen presentation and costimulatory molecules on *Arid1a*-deficient TAMs led us to further characterize CD8+ T cells in the tumor microenvironment in *Arid1a^f/f^;LysM^Cre^* mice. Flow cytometry analysis revealed over double the number of CD8+ infiltrating T cells in the tumors of untreated *Arid1a^f/f^;LysM^Cre^* hosts versus control hosts (Fig. 6A). As expected, CD8+ T cell infiltration was increased with anti-PD-L1 treatment in both genotypes (Fig. 6A). In contrast, the proportion of IFN-γ+GZMB+ CD8+ T cells was significantly higher in *Arid1a^f/f^;LysM^Cre^*versus control hosts in both untreated and anti-PD-L1 treated conditions (Fig. 6B-C), indicating increased CD8+ T cell activation in the presence of *Arid1a*KO TAMs. There were no changes in the proportion of PD-1+ cells, nor exhaustion subsets defined by SLAMF6 and TIM-3 expression between genotypes within each treatment group (Supplementary Fig. 11A-C).

**Fig. 6:**
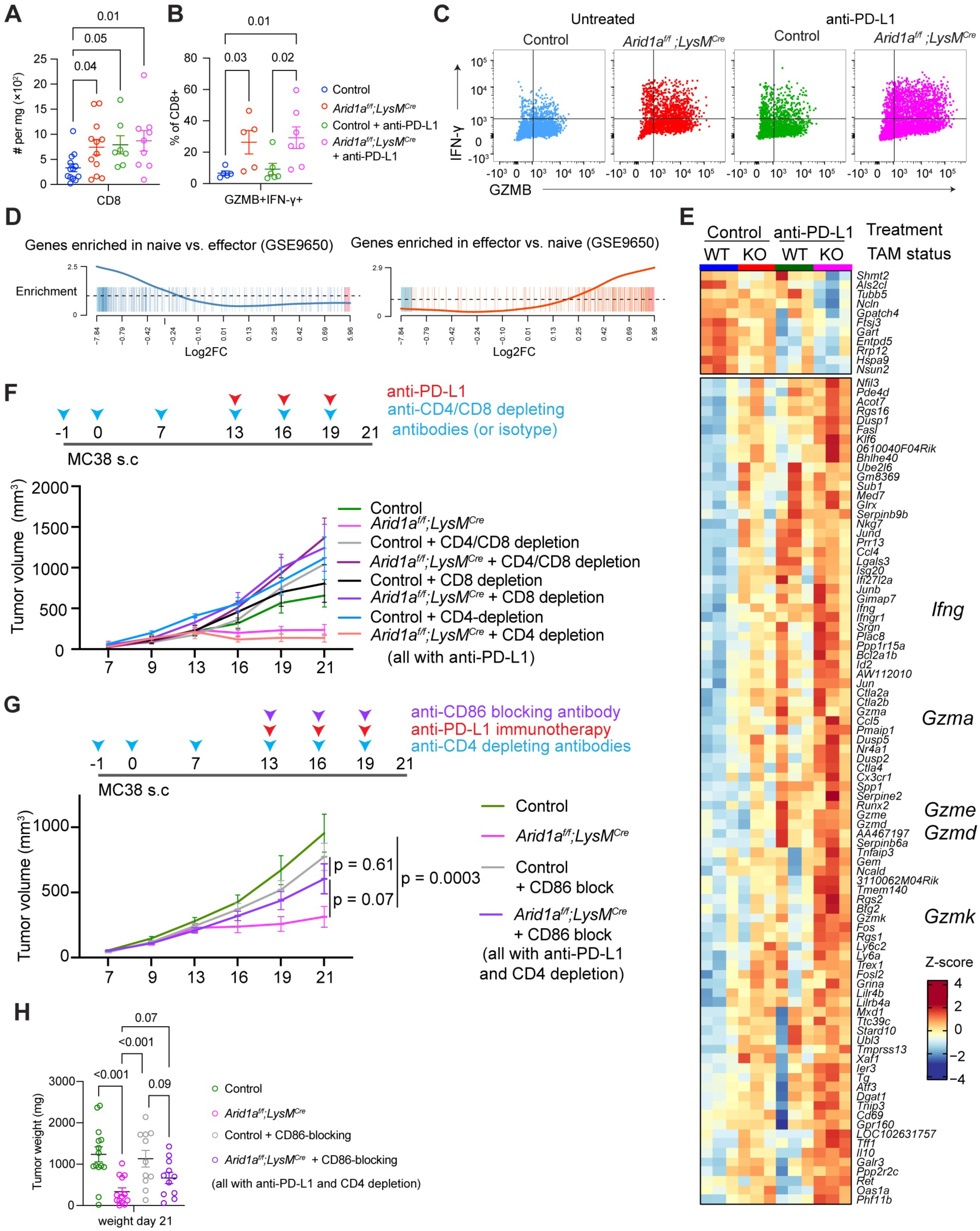
Loss of myeloid-specific ARID1A leads to increased CD8+ T cell activation and increased tumor-control dependent on CD8+ T cells and CD86 costimulation. A. Quantification of CD8+ T cell infiltration per mg tumor in mice of the genotypes indicated by legend in (B). B. Proportion of intratumoral CD8+ T cells expressing both GZMB and IFN-γ in mice of the indicated genotype. C. Representative cytokine staining for data shown in (B) in CD8+ T cells from tumors implanted in mice of the indicated genotypes. D. Barcode plots showing downregulation of naïve vs effector CD8+ T cell gene signatures indicating and upregulation of effector vs naïve genes in CD8+ T cells from *Arid1a^f/f^;LysM^Cre^* mice compared to controls (from untreated/isotype-only tumors). E. Genes significantly differentially expressed (adjP < 0.05) in CD8+ T cells from tumors in anti-PD-L1 treated *Arid1a^f/f^;LysM^Cre^* mice compared to CD8+ T cells from untreated (isotype-only) controls. Row z score indicates intermediate expression in CD8+ T cells from untreated *Arid1a^f/f^;LysM^Cre^* mice and from anti-PD-L1 treated controls. F. Tumor volume of mice of indicated genotype subcutaneously implanted MC38 tumors and treated with anti-PD-L1 plus either CD4, CD8 or combined depletion compared to isotype-only treated controls. G. MC38 tumor volume in mice of indicated genotypes with indicated treatments, showing CD86-blocking antibody partially reverses the increased response to anti-PD-L1 caused by ARID1A-myeloid lineage deletion (depletion of CD4 throughout the duration of the experiment to focus on CD8+ T cells and avoid confounding effects from CTLA-4 on Treg cells). H. MC38 tumor weights at endpoint of indicated genotypes and treatment. Data in (A,) (B), (H) were analyzed by one-way ANOVA. (H) was square-root transformed prior to statistical analysis to more closely fit normal distribution. Longitudinal tumor-growth in (F) and (G) analyzed using a linear mixed model with breakpoint at day 13 and assessed by type II ANOVA. Sequencing data analyzed as described in methods. Data presented as mean ± SEM. Where shown, each datapoint represents a biological replicate.

To further explore the phenotype induced in CD8+ T cells due to ARID1A loss in myeloid cells, we performed RNA-sequencing on sorted CD8+ T cells from MC38 tumors in control and *Arid1a^f/f^;LysM^Cre^*mice treated with or without anti-PD-L1. Probing for CD8+-specific genesets in the Immunologic Gene Signatures from MsigDB, we identified pathways significantly different between genotypes within each condition (Supplementary Table 4). Among the top pathways enriched in CD8+ T cells from untreated *Arid1a^f/f^;LysM^Cre^*versus controls were gene signatures comparing effector and naïve CD8+ T cells (Fig. 6D), consistent with increased CD8+ T cell activation. Indeed, we observed increased expression of many genes involved in CD8+ T cell function in *Arid1a^f/f^;LysM^Cre^* mice relative to untreated controls, including *Ifng* and several granzyme genes (Fig. 6E), which was even more pronounced with anti-PD-L1 treatment. This indicates that loss of ARID1A in myeloid cells influences the transcriptome and function of intratumoral CD8+ T cells.

## CD8+ T cells are required for the increased anti-PD-L1 response in tumors with myeloid-specific *Arid1a* deletion

Given increased activation of CD8+ T cells, we tested the functional requirement of T cells in the increased response to anti-PD-L1 observed in *Arid1a^f/f^;LysM^Cre^* mice. We determined that the phenotype was dependent on CD8+ cells but independent of CD4+ cells, as antibody-based depletion of CD8+ cells or combined CD4+/CD8+ removed tumor control, while tumor control remained enhanced in *Arid1a^f/f^;LysM^Cre^*mice with depletion of CD4+ cells alone (Fig. 6F, Supplementary Fig. 11D-E).

We next sought to identify specific molecules mediating interactions between CD8+ T cells and *Arid1a*KO TAMs that contribute to the elevated immunotherapy response. We hypothesized CD86 could contribute to the enhanced immunotherapy response, given its upregulation in *Arid1a*KO TAMs at both the transcript and protein level. Notably, blockade of both CD86 and CD80 eliminates the efficacy of anti-PD-L1 immunotherapy in the CT26 murine colorectal cancer model^92^, indicating costimulation is required for immune checkpoint blockade response. Using a similar strategy to test the effect of CD86 costimulation^92^, we found that the efficacy of the anti-PD-L1 response in the *Arid1a^f/f^;LysM^Cre^* hosts was reduced with CD86 blockade (Fig. 6G-H). This effect was specific to the context of *Arid1a* deletion as there was no effect of CD86 blockade in control anti-PD-L1 treated tumors. We confirmed CD86 was efficiently blocked and there was no significant change in CD80 expression (Supplementary Fig. 11F). These data demonstrate the requirement for costimulation via CD86 in the immunotherapy response in tumors lacking myeloid ARID1A.

## Discussion

Here, we identify the ARID1A-containing SWI/SNF complex as a key regulator of TAMs. ARID1A loss resulted in reduced enhancer accessibility and expression of tumor-promoting genes such as *Spp1*, together with increased promoter accessibility and expression of ISGs, effects that were phenocopied with pharmacological SWI/SNF ATPase inhibition. CD8+ T cells were more highly infiltrated and activated in *Arid1a^f/f^;LysM^Cre^*tumors and were required for the increased efficacy of immune checkpoint blockade in hosts lacking myeloid ARID1A. Together, these findings support a model in which ARID1A-containing cBAF helps maintain the regulatory landscape that sustains tumor-promoting TAM states while constraining the interferon response in macrophages.

While TAMs have typically been considered immunosuppressive, it is increasingly clear that if appropriately activated, TAMs can have anti-tumor potential particularly in combination with T cell-mediated immunotherapies. A dominant transcriptional signature impacted following loss of ARID1A in TAMs was upregulation of ISGs, including *Cd86.* Our data point to cell intrinsic and extrinsic contributions to ISG upregulation; namely, cell-intrinsic changes prime the response to exogenous IFNγ released by CD8+ T cells. Chronic antigen presentation by macrophages in the tumor-microenvironment contributes to T cell exhaustion^93–95^, but can also play a role in tumor clearance^11–14^. Our work highlights the importance of costimulation from CD86 in the productive anti-tumor immune response driven by myeloid-specific ARID1A loss. Notably, dual blockade of CD80 in combination with CD86 eliminates the efficacy of anti-PD-L1 immunotherapy in wildtype mice^92^. In contrast, CD86 blockade alone (with CD80 intact) selectively reduced immunotherapy efficacy in *Arid1a^f/f^;LysM^Cre^* but not control mice. This context-specific effect may reflect altered stoichiometry of B7 (CD80/CD86) molecules^96,97^ on *Arid1a*KO TAMs which would alter interactions with CD28 activating and CTLA-4 inhibitory receptors on CD8+ T cells. In addition, we expect the upregulation of MHC class I^12^ and other ISGs including SLAMF7^98^ could also contribute to the beneficial TAM:CD8 interactions in the tumor microenvironment.

At the molecular level, transcriptional reprogramming in *Arid1aKO* TAMs is accompanied by changes in the chromatin accessibility landscape. We paired our chromatin accessibility analysis with CUT&RUN data to assess key histone modifications associated with gene regulatory elements to build a genome-scale enhancer map of TAMs. The primary role of the SWI/SNF complex is in opening chromatin^99–101^, and consistent with this function, the sites most vulnerable to loss of ARID1A were poised and active enhancers, enriched for known binding partners of the SWI/SNF complex, including the AP-1 family^84–86^ and PU.1^38–40^, suggesting that ARID1A may be required to initiate and maintain binding of these factors. In the absence of SWI/SNF-dependent remodeling, TAMs fail to adopt subtype-specific accessibility landscapes associated pro-tumor TAM subtypes, including the SPP1+ and MRC1+ expressing subtypes. Indeed, we observed a reduction in *Spp1* in *Arid1a^f/f^;LysM^Cre^* and SWI/SNF inhibitor treated mice, associated with the loss of accessibility at enhancers of *Spp1*. As SPP1 secretion by TAMs was described recently as a key predictor of outcomes in solid tumors^21^, the loss of this pro-tumor function may contribute to the tumor control we observe in *Arid1a^f/f^;LysM^Cre^*mice.

In addition to loss of TAM subtype-specific expression, we observed an increase in ISGs across all TAM subtypes and in *Arid1a*-deficient or BRDK98-treated BMDMs. This is in contrast to our previously reported findings in which acute SWI/SNF inhibition blocked induction of ISGs and inflammatory genes in response to endotoxin^46,47^, and may reflect distinct biology arising from long-term (days) versus acute (hours) disruption of cBAF activity as well as homeostatic versus stimulation-induced responses. Mechanistically, ISG upregulation in *Arid1a*KO TAMs is independent of IFNAR and other secreted factors and associated with increased chromatin accessibility, unlike our previous observations of ISG upregulation in *Arid1a*KO cancer cells, which we found was independent of accessibility changes and driven by STING-dependent IFNAR signaling^65^. Increased gene expression/accessibility upon loss of SWI/SNF function has been ascribed to indirect mechanisms, including ARID1A-dependent recruitment of histone deacetylases^102,103^, redistribution of EP300^101^, Polycomb antagonism^104^ and increased binding of EP400 and SNF2H^105,106^. In *Arid1aKO* TAMs, IRF motifs were enriched in increased accessible sites, suggesting increased transcription factor binding may contribute to increased ISG expression. Interestingly, recent work has indicated a role of IRF1/IRF2 crosstalk in IFNAR-independent regulation of ISGs in macrophages, indicating an altered balance of IRF1/IRF2 binding could be mediating the IFNAR-independent upregulation of ISGs in the context of *Arid1a-*deletion^107^.

SWI/SNF ATPase inhibition was more effective at slowing tumor growth in the presence of anti-PD-L1 than myeloid ARID1A genetic deletion, likely due to effects on tumor and immune cells. Indeed, we report here an analysis of the entire microenvironment through paired chromatin accessibility and gene expression profiling revealing changes in tumor cells, CD8+ T cells and macrophages. We previously showed that ARID1A mutation in tumor cells can boost anti-tumor immunity^65^, suggesting direct effects of FHD-286/BRM014 on tumor cells likely contribute to immunotherapy efficacy. FHD-286/BRM014 also increased CD8+ T cell cytokine secretion and reduced TOX expression, consistent with several studies reporting that SWI/SNF complex disruption enhances CD8+ T cell effector function and reduces exhaustion^80,108–111^. Additionally, we speculate that systemic SWI/SNF inhibition might affect tumors through myeloid output from the bone marrow^112^, an effect we were not able to capture in *Arid1a^f/f^;LysM^Cre^*mice due to the persistence of ARID1A protein in myeloid precursors and monocytes. Finally, FHD-286/BRM014 disrupt the catalytic activity of all SWI/SNF complexes, and are thus molecularly distinct from loss of ARID1A, which we would predict would affect the catalytic and non-catalytic functions of the ARID1A-containing cBAF complex, but not ARID1B-containing cBAF complexes, PBAF or ncBAF.

SWI/SNF complexes are attractive therapeutic targets for synthetic lethal or combination therapy approaches. We found that SWI/SNF inhibition reprograms the tumor microenvironment, leading to increased immune checkpoint blockade sensitivity. Our genetic studies in *Arid1a^f/f^;LysM^Cre^*tumors clarify a role for the ARID1A-containing cBAF complex in TAMs, driven at the molecular level by changes in chromatin accessibility and gene expression. We demonstrate that ARID1A deletion in TAMs is sufficient to enhance response to immune checkpoint blockade and we identified CD8+ T cells and CD86 as key effectors of this response. These data suggest that effects on the tumor microenvironment, including TAMs, should be considered in the application of SWI/SNF targeted therapies and raise the possibility of targeting the SWI/SNF complex in macrophage reprogramming for example through directed delivery or CAR-macrophage engineering^113^.

## Acknowledgments

We thank the Hargreaves lab and Salk Institute NOMIS Center for their valuable discussion of this work, including Kaech lab members for advice and resources including the MC38 cell line and P14 mice. We thank Rose Marie Gonzales and Caitlin Chambers for excellent technical support. This work was supported by the following Salk Institute core facilities: Flow Cytometry, Next Generation Sequencing, and Integrative Bioinformatics with support from NIH-NCI CCSG P30 014195 and Shared Instrumentation Grant S10-OD023689 (Aria Fusion cell sorter). This work was supported by R01 CA285867 (D.C.H., E.C.D.), R01 AI066232 (S.M.K., D.C.H.) and the Pew-Stewart Scholars for Cancer Research Award to D.C.H. H.M.M was supported by the Catharina Foundation Fellowship, a Tang Prize Fellowship, American Australian Association scholarship, Salk Women and Science Special Award, and a Cancer Research Institute/Merck Postdoctoral Fellowship (CRI Award #4045). H.R.M. was supported by the NIH under the Molecular Biological Approaches to Endocrinology Institutional National Research Service Award Training Grant (T32DK007541-33) and the Ruth L. Kirschstein National Research Service Award Individual Predoctoral Fellowship to Promote Diversity in Health-Related Research from the NHLBI (F31HL167460). S.A.R and B.Y.C. were supported by NIH grant 5T32GM133351.

## AUTHOR CONTRIBUTIONS

Conceptualization, HMM, DCH; Formal Analysis, HMM, KGE, MJB, BYC; Funding Acquisition, HMM, DCH, SMK, DR; Investigation, HMM, KMN, QVM, SAR, BTS, MJB, JCB, HRM, IC; Supervision, HMM, DCH, DR; Visualization, HMM; Resources, KK, ECD, SMK; Writing – Original Draft, HMM, DCH; Writing – Review & Editing, HMM, DCH.

## Methods

### Genetic mouse models

*Arid1a^f/f^* mice^70^ (Magnuson lab) were backcrossed to C57BL/6J mice (Jackson lab strain #000664) for 3 generations using a marker assisted selection (i.e. “speed congenic”) approach. Mouse genomes were assessed at the DartMouse™ Speed Congenic Core Facility at the Geisel School of Medicine at Dartmouth. DartMouse uses the Illumina, Inc. (San Diego, CA) Infinium Genotyping Assay to interrogate a custom panel of 5307 SNPs spread throughout the genome. The raw SNP data were analyzed using DartMouse’s SNaP-Map™ and Map-Synth™ software, allowing the determination for each mouse of the genetic background at each SNP location. Genetic background at the final back-cross generation was determined to be 99.4% for the C57BL/6 background. Further backcrossing was performed to Jackson lab mouse strains on C57BL/6J background harboring Cre alleles (*LysM^Cre^ #*004781^71^, *LysM^CreERT2^* JAX#032291^72^, *Cx3cr1^Cre^* JAX#025524^114^, *Itgax^Cre^* JAX#008068^73^, *UBC^CreERT2^* JAX#007001^115^ and regularly refreshed by additional crosses to C57BL/6J (JAX#000664). Genotyping was performed through Transnetyx. A combination of controls were used to control for both the *Arid1a* floxed alelle and Cre alleles, and Cre-alleles were heterozygous in most cases (information on exact genotypes for each experiment available on request). Both female and male mice were used depending on tumor type (see tumor studies section), and mice used in experiments were aged between 8 to 22 weeks, using littermate controls where possible, or age-matched otherwise. For BMDM co-culture studies, CD45.1+ mice were purchased from the Jackson lab (JAX#002014). P14 mice (Kaech lab) have been previously described^116^. Animals were housed in specific-pathogen-free facilities at the Salk Institute, housed on a 12-h light-dark cycle, at an ambient temperature of 23 °C, and 30-70% humidity. Water and food were provided ad libitum. All experimental studies were approved and performed in accordance with guidelines and regulations implemented by the Animal Resources Department at the Salk Institute for Biological Studies and Institutional Animal Care and Use Committee (IACUC).

## Tumor cell lines

Tumor cells (MC38; sourced from the Kaech laboratory, Salk, or LLC; a gift from the Varner laboratory, UCSD) were cultured in DMEM (Corning, 10-13-CV), 10% FBS (Corning, Cat#35-010-CV), and 1% pen/strep (Gibco, Cat#11510-122), with the addition of non-essential amino acids (Gibco, Cat#11140-050). PyMT-ChOVA-mCherry cell line^75^, sourced from the Kersten laboratory, SPBD, were cultured in DMEM (Corning, Cat#10-13-CV), 10% FBS (Corning, Cat#35-010-CV), and 1% pen/strep (Gibco, Cat#11140050). Cell lines were tested as mycoplasma negative.

## Murine tumor studies and treatments

All procedures and treatments were performed under isoflurane anesthesia. For subcutaneous implantation, tumor cell lines were detached by trypsin (0.05% for MC38, Gibco, Cat#25300-054 and 0.25% for LLC, Gibco, Cat#25300-056), washed and resuspended at a concentration of 2.5 million cells per mL in PBS (Salk). Cells (0.5 million in 200 µL) were injected using a 27G needle (BD, Cat#305109) into mice of indicated genotypes, using female mice for MC38 and male mice for LLC experiments.

For mammary tumor studies, PyMT-ChOVA-mCherry cells^75^ were detached using trypsin (0.05%, Gibco Cat#25300-054) washed in PBS and resuspended in HBSS (Gibco, Cat#14175-095) with 2% FBS. A total of 2.5 x 10⁵ cells (in 7 µL) were mixed with 1 µL of trypan blue (0.05%, Sigma, Cat#T8154) to facilitate visualization of the injection site, and 8 µL of growth factor reduced Matrigel (Corning, Cat#356231). Cell suspensions were orthotopically implanted into the fourth mammary fat pad of 9- to 12- week-old female mice using pre-cooled 0.3 cc, 31G ultrafine insulin syringes (BD, Cat#328438). Skin flaps were sutured and stapled following transplantation.

Anti-PD-L1 immunotherapy (200 µg of clone 10F.9G2, Bio X Cell, Cat#BE101) or isotype (rat IgG2b clone LTF-2, Bio X Cell, Cat#BE0090) was provided intraperitoneally at day 13, 16 and 19 for subcutaneous MC38 experiments and day 13 and 16 for mammary tumor experiments. Tamoxifen (Sigma, 10540-29-1) was dissolved in peanut oil (Sigma, Cat#8002-03-7) at a concentration of 20 mg/mL and administered intraperitoneally at a dose of 2 mg/mouse for 5 consecutive days (Supplementary Fig. 6C). FHD-286 (MedChemExpress, Cat#HY-144835) was administered once daily (Fig. 1A) by oral gavage at a dose of 1.5 mg/kg as described^56^ in a solution of 20% HP-β-CD Water (Thermo Fisher Scientific Cat#10977023) in HyPure Molecular Biology Grade Water (Hyclone, Cat#SH30523.02), used as the vehicle. BRM014 (MedChemExpress, Cat#HY-119374) was administered once daily (Supplementary Fig. 2A) by oral gavage at 20 mg/kg as described^37^ in a solution of 10% DMSO and 1% methylcellulose, which was used as the vehicle. For T cell depletion studies, 200 µg of the following antibodies were used with dosing schedule as in Fig. 6F: anti-mouse CD4 antibody (clone GK1.5, Leinco, Cat#C1333), anti-mouse CD8a (clone 2.43, Leinco, Cat#C380), rat IgG2b isotype control (clone 1-2 Leinco, Cat#I-1034) and/or rat IgG2a isotype control (clone 1-1, Leinco, Cat#I-1177). For CD86 blocking studies, 200 µg of anti-CD86 (clone GL1, Leinco, Cat#C2158) was used (Fig. 6G), performed in the context of CD4 depletion as previously described^92^.

Tumor volumes were calculated using digital caliper measurements where volume = 0.5 x length x width x width. IACUC-defined endpoints were when tumors reached 20 mm in diameter or if ulcerated. Statistical analysis for tumor growth was performed using linear mixed models fit using the TumGrowth package^117^, using type II ANOVA with time and treatment or genotype as independent variables. Mice that had a tumor that reached over 90 mm^3^ by first treatment were enrolled in the SWI/SNF inhibitor studies, and constitutive *Arid1a* deletion studies, and mice that reached over 60 mm^3^ at first treatment were included for tamoxifen-inducible experiments. Mice were allocated into treatment group following ranking based on caliper measurement to ensure comparable distribution of tumor sizes across treatment.

## CellTiter-Glo assay

MC38 cells were plated at a density of 5000 cells per well of a 96-well flat-bottom TC-treated plate (Genesee Scientific, Cat#25-221) in DMEM + 10% FBS + pen/strep + NEAA. After settling overnight, cells were treated with FHD-286 (MedChemExpress, Cat#HY-144835) dissolved in DMSO at a final dose of 10 nM or 100 nM or with doxorubicin (Cayman, Cat#25316-40-9) dissolved in DMSO at a final concentration of 1 µM as a positive control. CellTiter-Glo Luminescent Cell Viability Assay (Promega, Cat#G7571) was performed according to the manufacturer’s instructions and luminescence acquired on TECAN Infinite M1000 Pro plate reader.

## BMDM cultures and P14 CD8+ T cell co-culture

Bone marrow was harvested by flushing the femurs of adult (2 to 4 month old) mice and following red blood cell removal (BioLegend, Cat#420301), cultured in RPMI-1640 (Gibco, Cat#11360070) supplemented with 30% filtered L929 conditioned media, 20% FBS, 1% HEPES (Gibco, Cat#15630-080), 1% Sodium Pyruvate (Gibco, Cat#11360-070), 1% Penicillin/Streptomycin (Gibco, Cat#15140122), and 1% non-essential amino acids (Gibco, Cat#11140050) in 37°C with 5% CO_2_. Cells were cultured on Petri dishes for the first 6 days, with an equal volume media added at day 4 to maintain macrophage differentiation. At day 6, cells were lifted using cold PBS and plated onto TC plates and left to settle overnight in a 50:50 mix of BMDM media and DMEM (Corning, Cat#10-013-CM) supplemented with 10% FBS and 1% Penicillin/Streptomycin (Gibco, Cat#15140122). Treatment for stimulation was initiated at day 7 and harvested using Accutase (Sigma, Cat#A6964a) and cold PBS and a cell lifter (Corning, Cat#3008) for flow cytometry on day 8.

Where indicated, BMDMs were treated/stimulated with IFN-γ (0.2 ng/mL, Peprotech, Cat#315-05), interferon-α/β receptor antibody (anti-IFNAR; Leinco, clone MAR1-5A3, Cat#I-401, 1 µg/mL), isotype antibody (mouse IgG1, Leinco, clone HKSP, Cat#I-536, 1 µg/mL), JAK1/2 inhibitor ruxolitinib (InvivoGen; Cat#TLRL-RUX, 0.5 µM), BRDK98^90^ (10 µM), or 4-hydroxytamoxifen (1 µM, Sigma Cat# 68392-35-8).

For co-culture experiments with P14 CD8+ T cells^116^, 5 x 10^6^ BMDMs in were incubated at 37°C for 1h in 1 mL media with 1 µg/mL GP33 peptide (GenScript, LCMV gp33–41, Cat#RP20257), with agitation every 15 minutes and washed three times prior to plating 1 x 10^5^ per well of a 12-well cell culture plate. P14 CD8+ T cells were isolated by negative selection from spleens that were mechanically dissociated by mashing through a 70 µm cell strainer (Falcon, Cat#352350) with the end of a 3 mL syringe (BD Medical, Cat#301073). Single cell suspensions were incubated in with biotinylated antibodies from Biolegend (anti-B220 clone RA3-6B2, anti-CD11c clone N418, anti-CD11b clone M1/70, anti-CD4 clone GK1.5, anti-CD49b clone DX5, anti-TCRγδ clone GL3, anti-Ter119 clone TER-119) in PBS + 1% rat serum (StemCell, Cat#13551) + 2% FBS + 1mM ETDA. After washing, cells were resuspended in 5% v/v MojoSort Streptavidin Nano beads (Biolgend, Cat#480016), incubated for 10 minutes and supernatants collected using a magnetic rack. Pre-activated CD8+ T cells were isolated the day prior to co-culture on plates pre-coated with goat anti-hamster IgG (30 µg/mL in 500 µL for a 24 well plate) to allow binding of anti-CD3 and anti-CD28. 1 x 10^6^ cells CD8+ T cells per well were incubated overnight in 1mL media (RPMI-1640 + 10%FBS + Pen/Strep + 50μM β-ME), supplemented with 2 µg/mL anti-CD3 (BD Cat#553057) and 0.5 µg/mL anti-CD28 (BD Cat# 553294) and 10 ng/mL recombinant human IL-2 (Peprotech, Cat# 200-02). 4 x 10^5^ pre-activated or freshly isolated P14 CD8+ T cells were added to a 12 well plate containing BMDMs in RPMI-1640 + 10%FBS + Pen/Strep + 50μM β-ME supplemented with 10 ng/mL recombinant human IL-2. After 72 h, non-adherent cells were collected by pipetting and adherent cells were lifted using Accutase (Sigma, Cat#A6964a), cold PBS and a cell lifter (Corning, Cat#3008) and processed for flow cytometry.

## Isolation of single-cell suspension from tissue and tumors for flow cytometry

For studies involving *Arid1a* genetic deletion, subcutaneous tumors were mechanically dissociated by chopping followed by enzymatic digestion as described^65^ with 0.05 mg/mL Liberase (Roche, Cat#5401020001) and 20 µg/mL DNase I (Roche, Cat#11284932001) in RPMI 1640 (Gibco, Cat#11875093), 20 mM Hepes (Gibco, Cat#1187509315630-080) and 1% pen/strep (Gibco, Cat#1187509315140-122) for 30 minutes at 37°C. For experiments involving BRM014 and FHD-286, the enzymatic digestion mix used gentle collagenase/hyaluronidase (StemCell, Cat#07919), 0.1 mg/mL DNase (StemCell, Cat#100-0762), 10% FBS, 25 mM HEPES in DMEM (Corning, Cat#10-13-CV) for 30 minutes at 37°C. Suspensions were mashed through a 70 µm cell strainer (Falcon, Cat#352350) with the end of a 3 mL syringe (BD Medical, Cat#301073).

For analysis macrophages of non-tumor bearing mice, spleens were dissociated using the same enzyme concentration as subcutaneous tumors, lung was enzymatically digested using the same protocol but with 0.5 mg/mL Liberase (Roche, Cat#5401020001). Liver was dissociated^118^ as described with 0.2 mg/mL Collagenase VIII (Sigma, Cat#C2139) and 20 µg/mL DNase I (Roche, Cat#11284932001) in 5% FBS in RPMI. Following 45 min incubation at 37°C, cells were centrifuged at 50 g for 3 min, followed by 400 g centrifugation of the aqueous phase prior to RBC lysis. Colon tissue was subject to epithelial segregation using 1 mM DTT, 2 mM EDTA, and 1% FBS in RPMI, followed by dissociation using 1 mg/mL collagenase VIII (Sigma C2139) and 20 µg/mL DNase I (Roche, 11284932001) and 1% FBS in RPMI. Peripheral blood was collected retro-orbitally into EDTA-coated tubes (BD, Cat#365974). Cells from all tissues were subject to RBC lysis (BioLegend, Cat#420301) and resuspended in PBS + 2% FBS.

## PMA/ionomycin re-stimulation

Single cell suspensions were plated onto a 96 well U-bottom plate (Genesee Scientific, Cat#25-221) in RPMI + 10% FBS + pen/strep and incubated at 37°C with 1 µM PMA (Selleck, Cat#S7791) and 1 µM ionomycin (StemCell, Cat#73724) for 4h, with addition of Brefeldin A (BioLegend, Cat#420601) after 15 minutes of initial stimulation. Cells were centrifuged (350 g) and processed for flow cytometry.

## Flow cytometry - analysis

Single cell suspensions were stained in a 96 well U-bottom plate (Genesee Scientific, Cat#25-221) with Zombie Red Fixable Viability dye (BioLegend, Cat#423110), followed by incubation with FC Block (Tonbo, Cat#70-016) in 2% FBS + 2 mM EDTA in PBS on ice for at least 10 minutes prior to incubation with fluorescent-conjugated antibodies (Supplementary Table 5).

Intracellular staining was achieved using the eBioscience™ Intracellular Fixation & Permeabilization Buffer Set (Invitrogen, Cat#88-8824-00), or for nuclear staining, the eBioscience Foxp3/Transcription Factor Staining Set (Thermo Fisher, Cat#00-5523-00). Intracellular/nuclear antibodies were fluorescent-conjugated or combined with a fluorescent-conjugated secondary antibody (listed in the Supplementary Table 5). Precision Count Beads (BioLegend, Cat#424902) were added to obtain absolute cell counts. UltraComp eBeads Compensation Beads (Thermo Fisher, Cat#01-2222-42) were used for single-stain compensation controls and data was acquired on a BD FACS Symphony A3 Cell Analyzer cytometer.

## Flow cytometry - sorting

Prior to staining, single cell suspensions were enriched for immune cells using CD45 TIL MicroBeads (Miltenyi, Cat#130-110-618) and LS columns (Miltenyi, Cat#130-042-401) for RNA-seq, scRNA-seq and Multiome experiments involving *Arid1a* genetic deletion, and H3K27me3 and H3K4me CUT&RUN experiments. A negative selection strategy was applied using biotinylated antibodies (anti-B220 clone RA3-6B2, anti-CD19 clone 6D5, anti-CD5 clone 53-7.3, anti-CD49b clone DX5), in 1% rat serum (StemCell, Cat#13551) with rapidSpheres (StemCell, Cat#19860) for H3K4me3 and H3K27ac CUT&RUN. A F4/80 positive selection strategy was employed (StemCell, Cat#100-0659) for the ATAC-seq experiment sort. No enrichment was performed prior to sorting for the FHD-286 or BRM014 experiments (RNA-seq or multiome) in order to include tumor cells.

Single cell suspensions of cells for sorting were incubated with FC Block (Tonbo, Cat#70-016), stained with fluorescent-conjugated antibodies (Supplementary Table 5) in polystyrene tubes (Falcon, Cat#352052) in 2% FBS + 2 mM EDTA in PBS, except for single-cell experiments where EDTA was excluded from all buffers from the staining step. After washing, cells were resuspended in a buffer containing 7-AAD viability dye (Biolegend, Cat#420404) in PBS+ 2% FBS + 2 mM EDTA + 25 mM HEPES + 100 µg/mL DNaseI (Sigma, Cat#10104159001) + 5 mM MgCl_2_ for all experiments except single cell experiments in which sort buffer contained no EDTA, DNaseI or MgCl_2_. Cells were filtered through cell strainer capped-tubes (Falcon, Cat#352235) prior to sorting on a BD ARIA Fusion with 100 µm nozzle. Gating strategies used are described in each section (bulk RNA-seq, ATAC-seq, CUT&RUN, scRNA, scMultiome).

## Bulk RNA-seq

### Bulk RNA sample preparation

For the bulk whole tumor FHD-286 vs vehicle samples, 30-50 mg of tissue from each tumor was isolated and transferred to an Eppendorf tube and frozen on dry ice. Samples were stored at -80°C until processing using a handheld homogenizer with plastic pestle for microtube (Cole-Parmer, Cat#UX-44468-19) following the addition of RNA lysis buffer (Zymo, Cat#R1060-1). RNA was isolated using a Zymo kit (Quick-RNA Miniprep Kit, Cat#R1055). RNA libraries were prepared using a TruSeq Stranded mRNA kit (Illumina) and sequenced on the NovaSeq X Plus instrument.

Gating strategy for sorted cells RNA-seq for the FHD-286 experiment were tumor cells: 7-AAD^neg^, CD45^neg^; TAMs: 7-AAD^neg^, CD45+, CD4^neg^, CD8^neg^, CD11b+, Ly6G^neg^, CD64+; CD8+ cells: 7-AAD^neg^, CD45+, CD11b^neg^, NK1.1^neg^, CD4^neg^, CD8+. For the *Arid1a*KO RNA-seq, two independent experiments each with 3 biological replicates per group were combined. For experiment 1, the gating strategy for TAMs was: 7-AAD^neg^, CD11b+, Ly6G^neg^, SiglecF^neg^, Ly6C^neg^, CD64+. For *Arid1a*KO experiment 2, TAMs were sorted using 7-AAD^neg^, CD45+, CD4^neg^, CD8^neg^, NK1.1^neg^, Ly6G^neg^, Ly6C^neg^, CD64+; and CD8+ cells were sorted using based on: 7-AAD^neg^, CD45+, CD11b^neg^, CD4^neg^, CD8+.

For sorted populations of tumor cells, TAMs and CD8+ cells, samples were sorted directly into lysis buffer (Qiagen RLT Plus with B-me added as per manufacturer’s instructions), diluted to a maximum of 0.5 x from the sort volume and frozen at -80°C. RNA was isolated using the manufacturer’s protocol (RNeasy Plus Micro Kit, Qiagen, Cat#74034), quality assessed using Tapestation (Agilent) and concentration through Qubit (ThermoFisher Scientific). cDNA was prepared using SMART-seq v4 kit (Takara, Cat#R400753) followed by Nextera XT (Illumina, Cat#FC-131-1096) for library preparation using 0.15 ng cDNA as input with Nextera XT Index Kit v2 Set A (Cat#FC-131-2001).

Libraries were pooled and diluted for paired-end 50bp sequencing on a NovaSeq 6000 instrument (*Arid1a*KO v control TAMs batch 1) or for single-end 100bp using the Nextseq 2000 instrument (FHD-286 bulk and sorted populations, *Arid1a*KO control TAM batch 2 and CD8 cells).

### Bulk RNA-seq data analysis

FASTQ reads were aligned to the mouse genome mm10 using STAR^119^, and raw count matrices generated using Homer script analyzeRepeats.pl^120^, counting reads in exons and one isoform per locus. Raw count matrices were imported into in R (v4.4.1) and using the edgeR (v4.2.0) pakage^121^, lowly expressed genes that failed to meet a minimum CPM threshold were filtered, TMM normalization^122^ was applied to each cell type followed by data transformation to log2(counts per million). Differential expression was assessed using linear modeling using Limma-Voom^123,124^, using quality weights^125^, and p-value adjustment using the Benjamini and Hochberg method.

Since two independent experiments were combined for the *Arid1a*KO versus control TAM RNA-seq, the linear model incorporated batch correction with experiment day as a variable. CD8-associated transcripts were detected in the first *Arid1a*KO versus control TAM RNA-seq experiment (which did not include CD8 negative selection step), likely reflecting doublets/ex vivo re-coupling^93^, or macrophage fragmentation^126^, thus a list of CD8-enriched genes were excluded prior to differential expression analysis.

Gene set enrichment analysis for MSigDB (v7.1) Hallmark collections^127^ or the immunologic signature gene sets^128^ was performed using CAMERA^129^ with the inter-gene correlation set to 0.01 using mouse genesets downloaded from https://bioinf.wehi.edu.au/MSigDB/, converting RefSeq annotation to ENTREZID using org.Mm.eg.db (v3.19.1). ROAST^130^ was used to compare specific data sets of interest^17,46,63,67^. Pearson’s correlation coefficients for macrophage reprogramming datasets^22,23,30,94^ were calculated in R, based on the log2FC correlation between conditions. Immune dictionary analysis^77^ was performed using the web software (available: www.immune-dictionary.org). Cell chat analysis^68,69^ was performed using CellChat (v2.1.2) in R, using a matrix of normalized logCPM data from bulk RNA-seq of sorted cell populations (tumor cells, TAMs and CD8+ cells), based on n = 6 biological replicates. A list of dendritic cell specific transcripts was filtered out of the CD8+ T cell matrix. Chord diagram visualizations were generated to exclude interactions contributing less than 0.01% and CD8:CD8 interactions. Barcode plots, volcano plots, correlation plots and heatmaps were generated using Limma (v3.60.3), gplots, EnhancedVolcano and ComplexHeatmap packages. All analyses were conducted in R (v4.4.1).

## Bulk ATAC-seq

### Bulk ATAC-seq sample preparation

TAMs (gated on 7-AAD^neg^, CD45+, CD8^neg^, CD4^neg^, CD19^neg^, CD11b+, Ly6G^neg^, Siglec^neg^, Ly6C^neg^, CD64+) from 3-4 biological replicates per group were sorted into a buffer containing RPMI-1640 (Gibco, Cat#11360070) supplemented with 20% FBS, 1% HEPES (Gibco, Cat#15630-080), 1% Sodium Pyruvate (Gibco, Cat#11360-070), 1% Penicillin/Streptomycin (Gibco, Cat#15140122), and 1% non-essential amino acids (Gibco, Cat#11140050). Omni-ATAC was performed as described^131^. Briefly, up to 50,000 cells were washed with cold PBS, collected by centrifugation then lysed in 50 µL ATAC resuspension buffer (10mM Tris-HCl, pH 7.4, 10mM NaCl, 3mM MgCl containing 0.1% NP40, 0.1% Tween-20, and 0.01% Digitonin in sterile water) on ice for 3 min. Following the addition of 1 mL wash buffer (10mM Tris-HCl, pH 7.4, 10mM NaCl, 3mM MgCl and 0.1% Tween-20 in sterile water), nuclei were pelleted and incubated in transposition mix (containing 2.5 µL Tn5 transposase and 25 µL TD buffer (Illumina, Cat#20034198) with 16.5 µL PBS, 0.5 µL 1% digitonin, 0.5 µL 10% Tween-20, 5 µL H_2_O) at 37°C for 30 mins, shaking. DNA was purified using a MinElute PCR Purification Kit (Qiagen, Cat#28006) and PCR amplified using indexed oligos and NEBNext High-Fidelity 2X PCR Master Mix (NEB, Cat#M0541L). qPCR was used to determine the number of amplification cycles. Amplified DNA was purified using MinElute (Qiagen, Cat#28006) and fragments between 100-1000bp were selected using AmpureXP beads (Beckman Coulter, Cat#A63882). Libraries were sequenced for 50bp paired-end reads on the NextSeq 2000 (Illumina).

### Bulk ATAC-seq data analysis

FASTQ reads were aligned to the mouse genome mm10 using STAR^119^. A common peak set was called using Homer findPeaks -style dnaseI set to 501 bp on merged reads from all samples. Following removal of peaks within regions mapping to mm10 blacklist regions^132^, differential accessibility between genotypes within each condition was called on the common peak set using the Homer^120^ pipeline getDifferentialPeaksReplicates.pl incorporating the default DEseq2^133^ for differential analysis, using -style dnase and -balanced. A consensus set of peaks sensitive to ARID1A loss was considered based on common log2FC between genotypes, with at least one condition meeting adjP < 0.05, and the p-value less than 0.05 in both conditions. For each biological replicate, a bedgraph file was generated using Homer’s makeUCSCfile, from which a representative BedGraph file for each group was created by averaging the normalized reads at each peak. Bedgraphs for each group were converted to bigwig for heatmap and profile plot visualization using deeptools^134^, and IGV^135^ for individual gene track visualization. Bedtools intersect function was used to annotate ATAC-seq peaks with ChIP-seq or CUT&RUN data to define ARID1A binding and regulatory regions.

## CUT&RUN

### CUT&RUN sample preparation

Following enrichment described in sorting procedures, TAMs (7-AAD^neg^, CD11b+, Ly6G^neg^, SiglecF^neg^, Ly6C^neg^, CD64+) were sorted into a buffer containing RPMI-1640 (Gibco, Cat#11360070) supplemented with 20% FBS, 1% HEPES (Gibco, 15630-080), 1% Sodium Pyruvate (Gibco, 11360-070), 1% Penicillin/Streptomycin (Gibco, Cat#15140122), and 1% non-essential amino acids (Gibco, Cat#11140050). H3K4me and H3K27me3 data were pooled from 5 biological replicates of wild-type control C57BL/6 mice and H3K4me3 and H3K27ac data were pooled from 4 biological replicates of each genotype (all MC38 tumors treated with 200 µg anti-PD-L1 on days 13, 16 and 21 post tumor implantation).

CUT&RUN was performed as described^136^ with the modification of using V-bottom plates (Thermo Scientific Nunc, Cat#249944) instead of magnetic beads to collect cells. Briefly, 70-150K cells were pelleted by centrifugation. Cells were washed in dig-wash buffer (20 mM HEPES, 150 mM NaCl, 0.05 mM spermidine, Roche Complete Protease Inhibitor EDTA-free, 0.005% digitonin) and incubated with antibodies overnight at 4°C (H3K4me3: Millipore, Cat#05-745R; H3K27ac: Millipore, Cat#MABE647; H3K27me3: Cell Signaling, Cat#C3B11; H3K4me: Abcam, Cat#Ab8895; or IgG control antibody: Cell Signaling Cat#3900S) in dig-wash buffer + 2 mM EDTA. Following 2 washes in dig-wash buffer, cells were incubated with pA/G-MNase (Epicypher, Cat#15-1116) for 1h at 4°C. Cells were washed twice in dig-wash buffer then resuspended in dig-wash buffer with 2 mM Cacl2 for 30 minutes on ice. Stop solution (340 mM NaCl, 20 mM EDTA, 4 mM EGTA, 0.005% digitonin, 100 µg/mL RNAse A, 50 µg/mL glycogen) was added and incubated 15 minutes at 37°C, with 0.5 ng of CUTANA E. coli Spike-in DNA (EpiCypher, Cat#18-1401) added per 1 x 10^5^ cells for H3K27ac and H3K4me3 assays. Supernatants were collected by centrifugation and DNA isolated using Qiagen MinElute. Libraries were prepared using a NEBNext Ultra II kit and sequenced for paired end sequencing on the NextSeq 2000 instrument.

### CUT&RUN analysis

FASTQ files were mapped using STAR^119^ and peaks were called using Homer findPeaks -style histone, against IgG control as background. Gene regulatory elements were defined based on histone regions called via Homer and based on control genotype data. Bigwig files were created from UCSC bedgraph file generated with Homer makeUCSCfile and visualized using deeptools^134^, and IGV^135^.

## 10X scRNA and multiome

### scRNA sample preparation

Single-cells (7-AAD^neg^ CD45+CD11b+) were collected in RPMI +10% FBS, washed once and resuspended in RPMI +10% FBS following centrifugation prior counting and loading on 10X Next GEM Chip K. Libraries were processed using Chromium Next GEM Single Cell 5’ Kit v2 (PN-1000265) following 10X protocol CG000331 Rev D. Libraries were sequenced for paired-end reads using the NextSeq 2000.

### scMultiome sample preparation

Sorted cells (TAMs for *Arid1a*KO and control samples gated on 7AAD^neg^, CD45+, CD4^neg^, CD8^neg^, CD19^neg^, CD11b+, Ly6G^neg^, SiglecF^neg^, Ly6C^neg^, CD64+ or all viable cells for BRM014 and vehicle samples gated on 7-AAD^neg^) were collected in PBS+ 0.04% BSA. Nuclei were isolated following the 10X demonstrated protocol CG000365, Rev C, with 0.5X diluted lysis buffer (diluted according to 10X protocol CG000366) for the BRM014 and vehicle samples. Nuclei were counted using a DeNovix cell counter prior to loading and multiome samples were processed using Chromium Next GEM Single Cell Multiome ATAC + Gene Expression Kit (PN-1000285) following 10X protocol CG000338 Rev F. *Arid1a*KO ATAC libraries were sequenced for paired-end reads on the NextSeq 2000, and *Arid1*aKO and ATAC GEX and BRM014 and vehicle ATAC and GEX libraries were sequenced for paired-end reads on the NovaSeq X plus.

### scRNA data analysis

Single-cell RNA was processed using cellranger-6.0.1. Downstream processing was performed using Seurat, filtering each sample to include cells with between 200-5500 features and less than 5% mitochondrial reads. Integrated data were further filtered for contaminating/doublet cells based on expression of lineages markers for T cells, B cells, NK cells and nonimmune cells to generate a TAM and monocyte Seurat object.

### scMultiome data analysis

For each sample, FASTQ files were processed using cellranger arc v7.1.0^137^. Filtered matrix files were processed using Seurat^138^ and Signac v1.15.0^139^. For BRM014 and vehicle datasets, samples were filtered to include nuclei with TSS enrichment > 1, 200-4000 features, 2000-100000 ATAC counts, and 500-10000 RNA counts. For the *Arid1a*KO and control datasets, nuclei in the bottom or top 15 percentile were removed for ATAC counts, RNA counts, and RNA features and filtered to include nuclei with a TSS enrichment > 2. RNA assays were normalized using SCTransform^140^, while ATAC assays were normalized using term frequency-inverse document frequency (TF-IDF) normalization^141^. Samples within the same experiment were integrated using Harmony v1.2.0^142^ for each assay. The join neighbor graph was computed using the harmony embeddings from each assay to create a joint UMAP visualization. For cell type level analysis, each cluster was determined by Seurat (resolution = 0.1). Cell types were evaluated using known markers for immune, tumor and endothelial cell types and using the markers for each cluster, as calculated by the FindAllMarkers function in Seurat.

ATAC counts for the common peak set were added from Fragment files using CreateChromatinAssay. Differential accessibility between conditions was calculated using pseudobulk aggregate counts for each sample/cell type combination with DEseq2 for differential expression analysis. Differentially expressed genes between conditions within each cell type were calculated using a pseudobulk approach (AggregateExpression in Seurat) followed by Limma/Voom pipeline for differential expression testing, with gene inclusion cutoff based on aggregate CPM. BigWig files for cell type/condition were generated from Signac Fragment files, by tiling the genome into 100 bp fixed-windows and normalizing read coverage on a per cell basis

Dataset integration of monocyte and TAM clusters from scRNA and scMultiome experiments:

Including only cells annotated as macrophages or monocytes, the RNA assays from 3 independent single-cell experiments (2 multiome experiments and one scRNA experiment) were filtered for genes found in all samples. Samples were integrated using Harmony v1.2.0^142^ and TAM/monocytes clusters were identified using a resolution of 0.1. Two minor clusters containing low RNA and feature counts were excluded resulting in 6 monocyte/macrophage clusters, annotated based on Seurat FindAllMarkers. Following integration and TAM/monocyte cluster identification, each dataset was then considered separately for comparisons between conditions/genotypes. For the multiome datasets, cells were matched based on common barcodes to incorporate the ATAC data. For the BRM014 versus vehicle dataset, differentially expressed genes between conditions were calculated using a pseudobulk approach (AggregateExpression in Seurat) followed by Limma/Voom pipeline for differential expression testing, with gene inclusion cutoff based on aggregate CPM. For the *Arid1a*KO versus control differential expression analysis, RNA from the scRNA was used given the higher capture of RNA from whole cells compared to nuclei, and assessed using Seurat FindMarkers, considering genes expressed in at least 10% of at least one cluster in the comparison. Pseudobulk ATAC-seq counts were generated for each biological replicate within macrophage clusters using Seurat AggregateExpression. Peaks were retained if they had nonzero counts in at least 10% of cells in 2 samples of one cluster. Differential accessibility between conditions or genotypes was tested using DEseq2 considering both treatment (anti-PD-L1 or untreated) and genotype in the model for *Arid1a*KO dataset. Peaks were annotated to their nearest transcription start sites (TSS) using the RefSeq database from the UCSC Genome Browser (assembly mm10). Genomic coordinates for all peaks were parsed into GRanges objects and the distance from each peak center to the nearest TSS was then calculated with the distanceToNearest function to identify promoters (within 1.5 kb of TSS) or enhancers (over 1.5kb from TSS). Annotation of distal chromatin-gene linkage was calculated using Signac LinkPeaks. ChromVAR^143^ was used to define motifs that separated clusters, using the JASPAR 2022 database^144^ and RunChromVAR in Signac. The top 10 motifs identified for each cluster were determined and one motif per family from this list was plotted for each condition. Density gene expression plots were generated using Nebulosa^145^. Trajectory analysis was performed using Slingshot^146^ (v2.1.4) on the scRNA data. Gene expression signatures were calculated using UCell^147^. Other visualizations were generated using DotPlot, CoveragePlot and FeaturePlot from Seurat and Signac.

## Data availability

Data from this study have been deposited at the GEO database under the accession number GSE324052.

---

**Supplementary Fig. 1:**
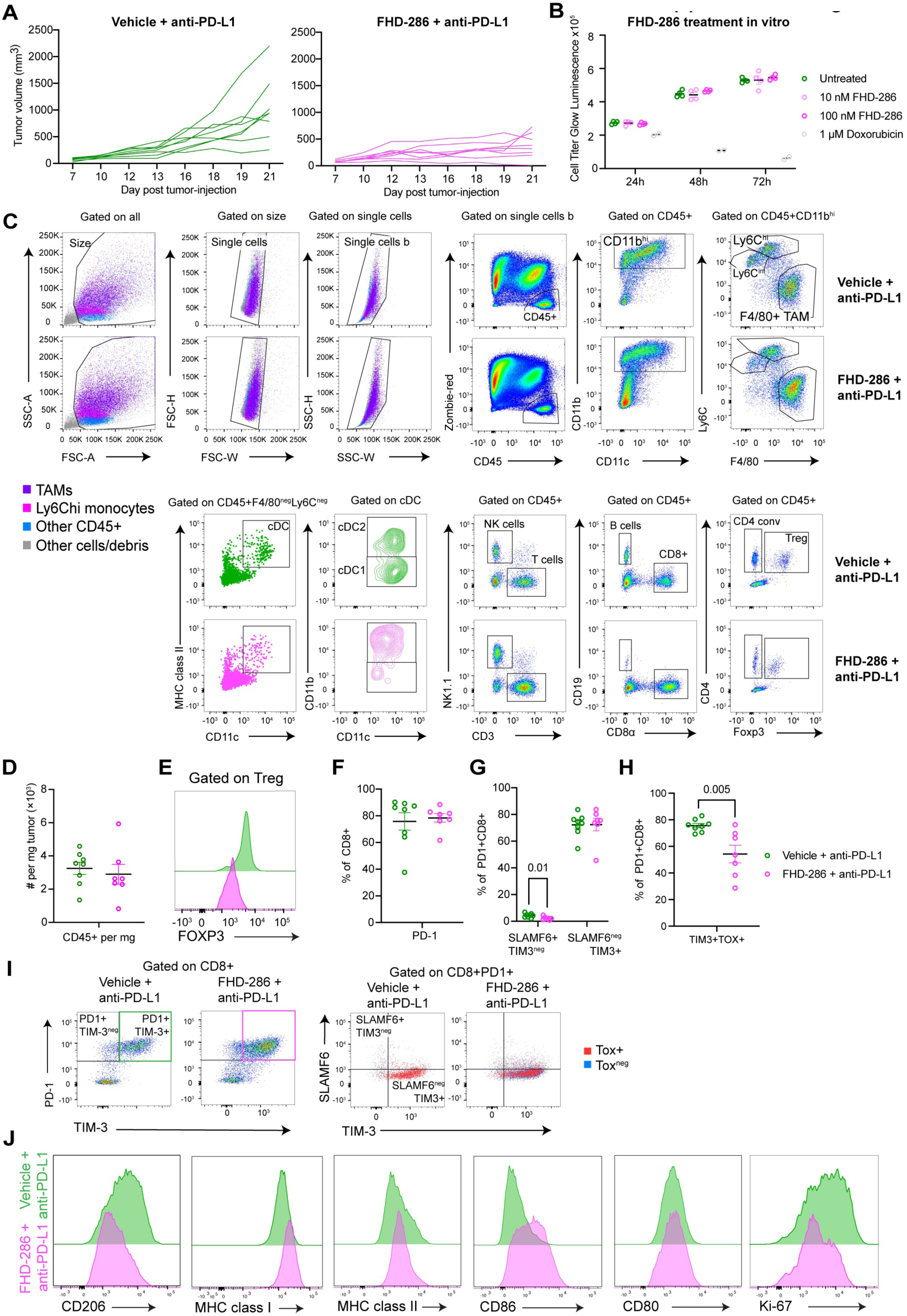
SWI/SNF complex inhibition with FHD-286 impacts tumor-infiltrating immune cells A. Growth curves showing individual biological replicates of data shown in Fig. 1A, treated with vehicle + anti-PD-L1 or FHD-286 SWI/SNF ATPase inhibitor + anti-PD-L1. B. *In vitro* growth of MC38 tumor cells in the presence of the indicated dose of FHD-286 compared to untreated cells (DMSO only) as assessed by Cell-Titer-Glo luminescence. C. Gating strategy used for the detection of intratumoral immune cell populations. D. Absolute quantification of intra-tumoral immune cell infiltration (CD45+) per mg of tumor. E. Representative histogram of Foxp3 protein expression gated on Treg cells (CD45+CD4+FOXP3+). F. Proportion of CD8+ T cells expressing PD-1. G. Proportion of CD8+PD-1+ T cells defined by the progenitor-exhaustion associated markers SLAMF6+TIM-3^neg^ and terminal-exhaustion associated markers SLAMF6^neg^TIM-3+. H. Proportion of CD8+PD-1+ T cells expressing the terminal-exhaustion associated markers TIM-3+ and TOX+. I. Representative plots showing PD-1 and TIM-3 expression and overlap of TOX+ cells on the SLAMF6 and TIM-3 quadrants. J. Representative histograms of cell surface markers on TAMs (CD45+CD11b+Ly6C^neg^F4/80+). Data analyzed by t-test between conditions.

**Supplementary Fig. 2:**
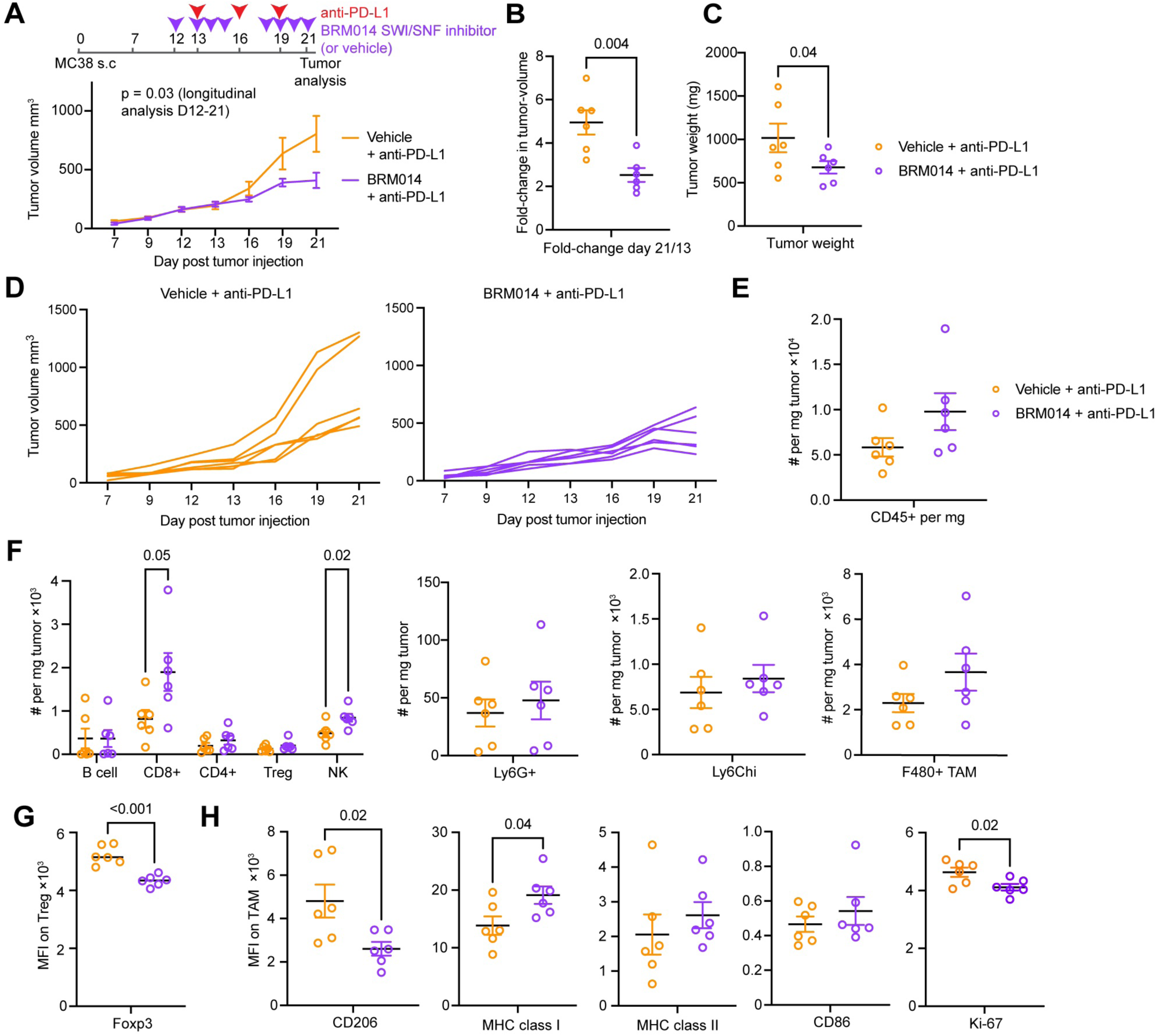
SWI/SNF inhibition using BRM014 causes similar effects on immunotherapy response and immune populations as FHD-286 treatment A. Schematic of experimental design and tumor growth in response to indicated treatments (20 mg/kg BRM014 delivered by oral gavage and 200 µg anti-PD-L1 intraperitoneal per dose). B. Fold change from first anti-PD-L1 treatment. C. Tumor weight for indicated conditions at day 21 (legend applies to all panels in this figure) D. Growth curves displayed to show individual biological replicates. E. Absolute quantification of all immune cells (CD45+) per mg tumor. F. Absolute quantification indicated immune populations per mg tumor. G. Cell surface expression of Foxp3 on Treg cells. H. Cell surface expression of indicated markers on TAMs. Data in A analyzed by a linear mixed model using the TumGrowth package with breakpoint at day 12 (treatment initiation) and subjected to a type II ANOVA and pairwise comparisons between conditions. Other data analyzed by t-test between conditions.

**Supplementary Fig. 3:**
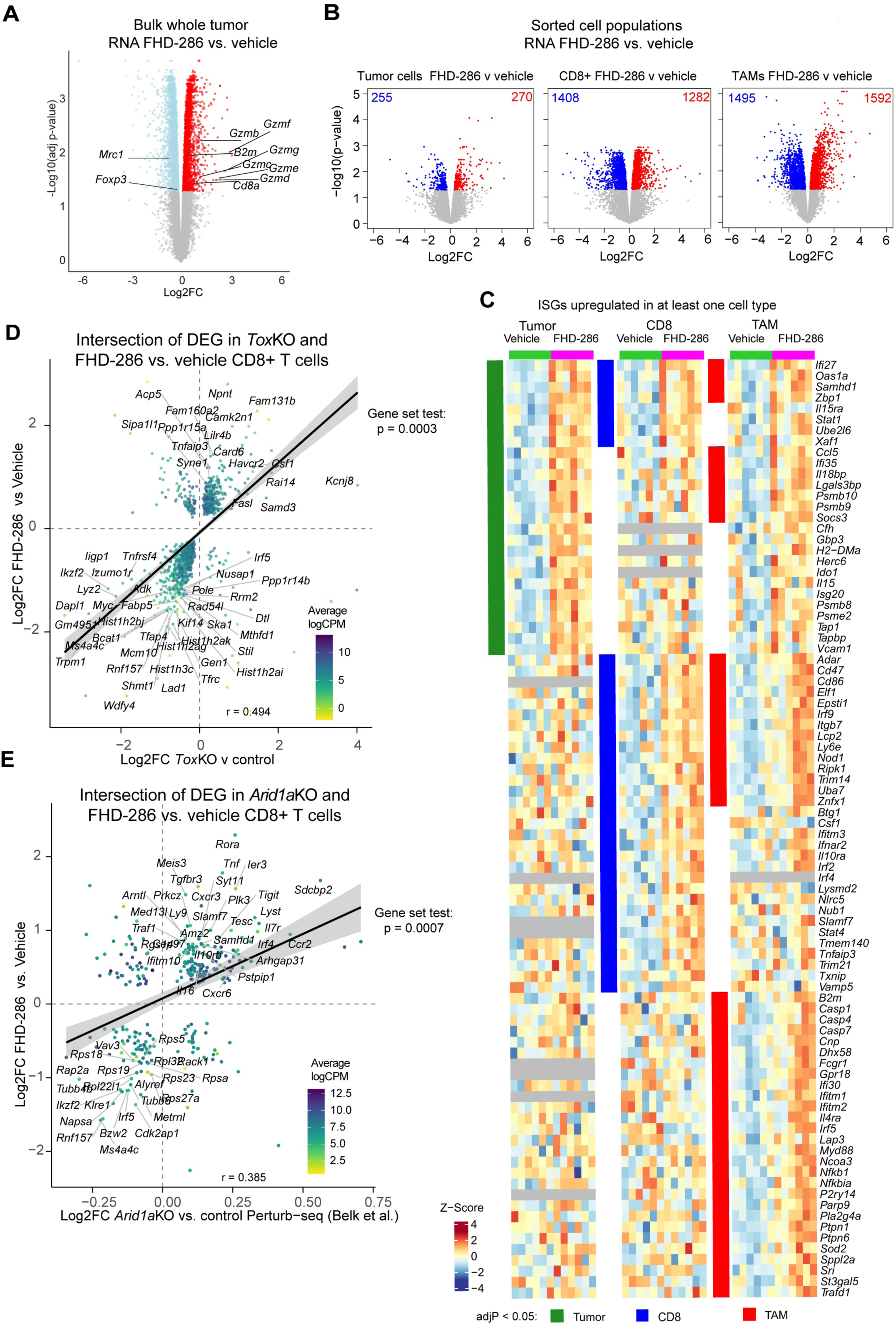
Transcriptional reprogramming of the tumor microenvironment by the SWI/SNF complex A. Volcano plot of bulk (whole tumor) RNA-seq of FHD-286 versus vehicle, highlighting selected immune genes of interest. B. Volcano plots highlighting significantly (adjP < 0.05) downregulated (blue) and upregulated (red) genes in indicated cell types from FHD-286 versus vehicle treated tumors (all anti-PD-L1 treated). C. Union of Hallmark interferon alpha and interferon gramma pathway genes that are upregulated in tumor cells, CD8+ T cells and TAMs after SWI/SNF inhibition (FHD-286 treatment). Significance within each cell type is indicated by color blocks as described in the legend. D. Correlation plot showing the intersection of differentially expressed genes in *Tox*KO CD8+ T cells (day 8 clone 13 LCMV infection from Khan et al. 2019^67^) and CD8+ T cells from FHD-286 versus vehicle (anti-PD-L1) treated tumors with selected genes highlighted. E. Correlation plot highlighting the intersection of differentially expressed genes in intratumoral *Arid1a*KO versus control CD8+ T cells (Perturb-seq dataset from Belk et al. 2022^63^) and CD8+ T cells from FHD-286 versus vehicle (anti-PD-L1) treated tumors.

**Supplementary Fig. 4:**
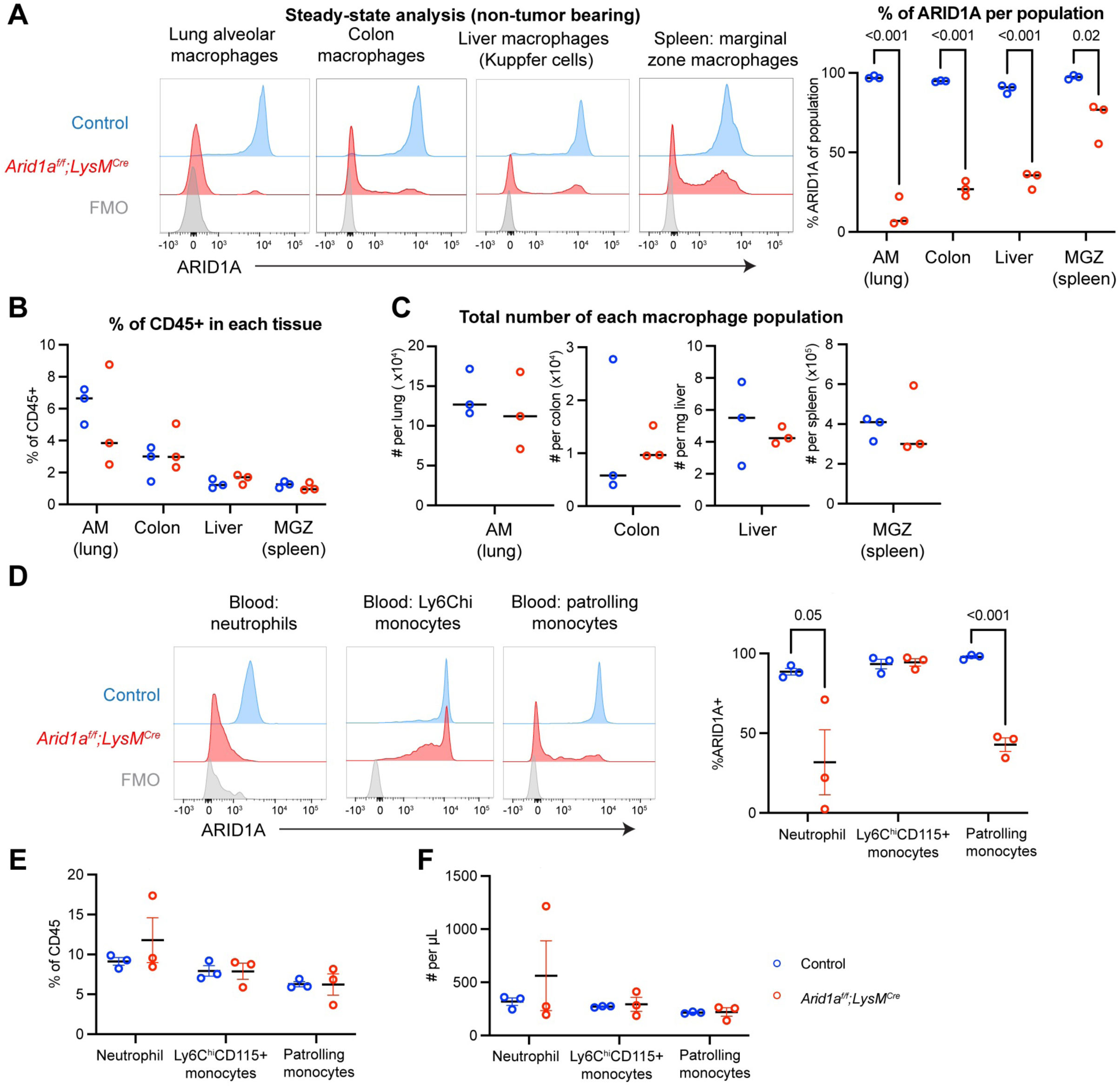
Characterization of myeloid-specific ARID1A deletion in non-tumor bearing mice A. ARID1A nuclear protein staining the indicated tissue-resident macrophage types of control or *Arid1a^f/f^;LysM^Cre^*genotype non-tumor bearing mice, showing representative gating on the left and quantification of the proportion of ARID1A+ cells on the right. B. Quantification of the proportion of indicated macrophage populations among CD45+ immune cells of control or *Arid1a^f/f^;LysM^Cre^*genotype mice. C. Quantification of the absolute number of tumor macrophages for the indicated tissues of control or *Arid1a^f/f^;LysM^Cre^* genotype mice. D. ARID1A nuclear protein staining in peripheral blood of non-tumor-bearing control or *Arid1a^f/f^;LysM^Cre^* genotype mice showing representative plots and quantification of ARID1A+ cells. E. Proportion of indicated myeloid populations among CD45+ cells in the peripheral blood of non- tumor-bearing control or *Arid1a^f/f^;LysM^Cre^* genotype mice. F. Absolute quantification of the indicated myeloid populations per µL in the peripheral blood of non- tumor-bearing control or *Arid1a^f/f^;LysM^Cre^* genotype mice. Data analyzed by t-test between conditions. Shared legend in panel F.

**Supplementary Fig. 5:**
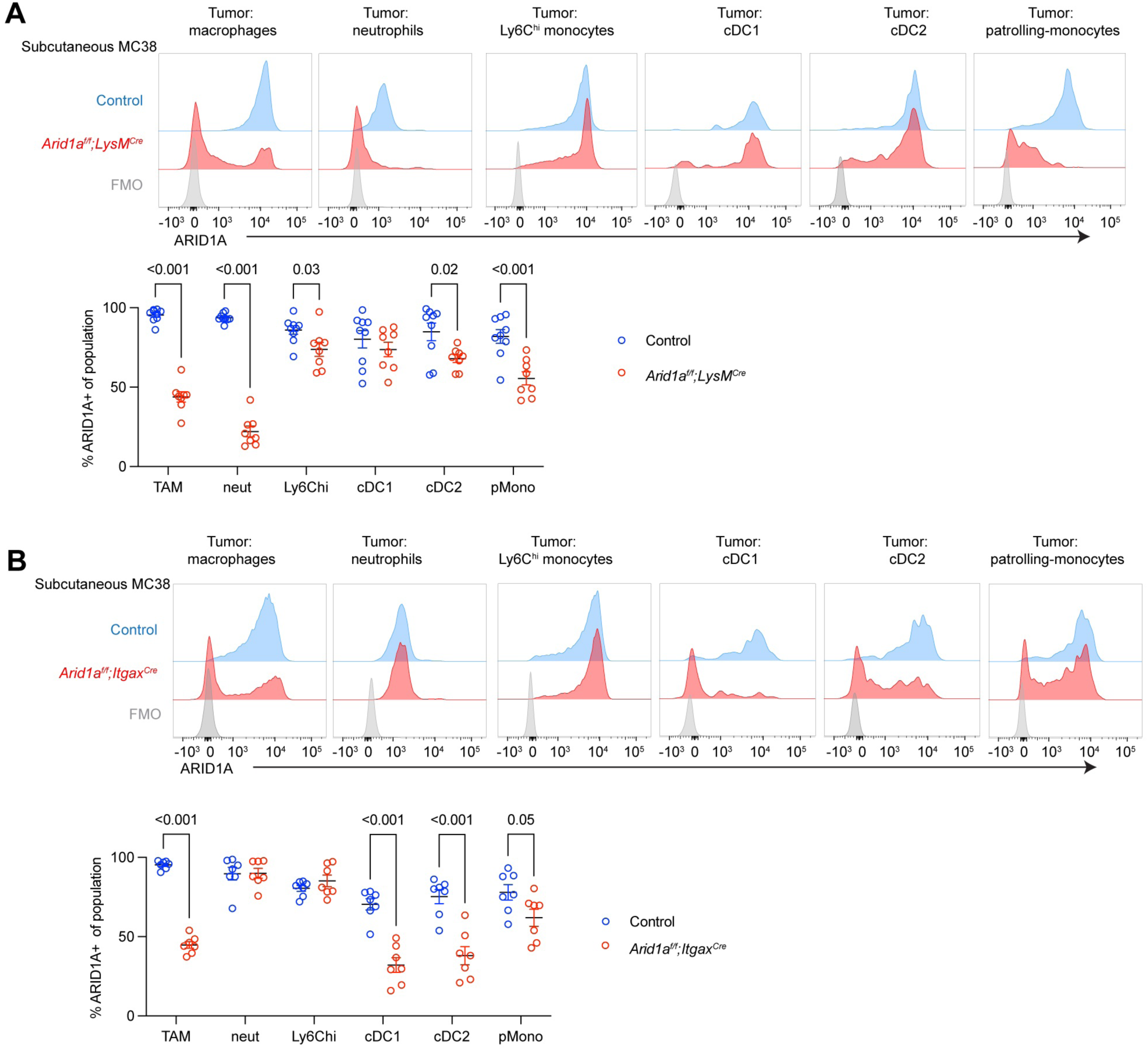
Characterization of ARID1A deletion efficiency in intratumoral myeloid cells A. ARID1A nuclear protein staining in the indicated intratumoral myeloid populations showing representative plots and quantification of ARID1A+ cells in control or *Arid1a^f/f^;LysM^Cre^*genotype mice. B. ARID1A nuclear protein staining in the indicated intratumoral myeloid populations showing representative plots and quantification of ARID1A+ cells in control or *Arid1a^f/f^;Itgax^Cre^*genotype mice. Data analyzed by t-test between conditions.

**Supplementary Fig. 6:**
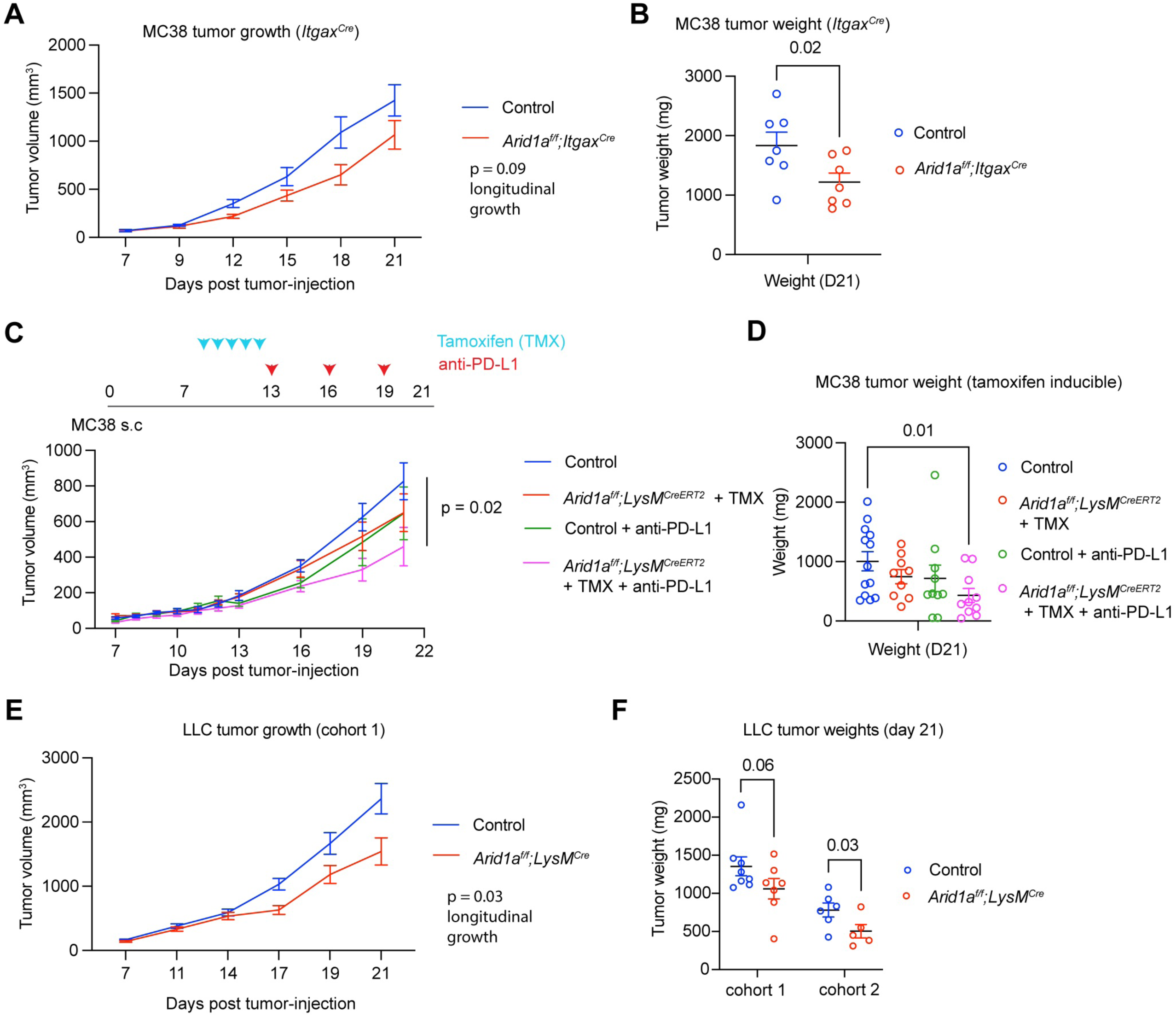
Additional tumor models with ARID1A deletion A. Tumor growth curve of MC38 tumors in control or *Arid1a^f/f^;Itgax^Cre^* genotype mice. B. Tumor weight in of day 21 MC38 tumors in control or *Arid1a^f/f^;Itgax^Cre^* genotype mice. C. Experimental design and tumor growth curves of MC38 tumors in tamoxifen-treated *Arid1a^f/f^;LysM^CreERT2^*compared to control mice with or without anti-PD-L1 treatment. P-value represents longitudinal analysis from day 13 to day 21. D. Tumor weight of MC38 tumors in tamoxifen-treated *Arid1a^f/f^;LysM^CreERT2^* compared to control mice with or without anti-PD-L1 treatment E. Tumor growth curves of Lewis Lung Carcinoma (LLC) tumor cells in control or *Arid1a^f/f^;LysM^Cre^*genotype mice showing one of two representative experiments. P-value represents results from longitudinal analysis. F. Tumor weight shown for two independent experiments (shown separately rather than pooled due to between-experiment variance). Data in A, C, E analyzed by a linear mixed model using the TumGrowth package and subjected to a type II ANOVA and pairwise comparisons between conditions. Other data analyzed by t-test between conditions. B and F analyzed by t-test and D analyzed by one-way ANOVA between conditions.

**Supplementary Fig. 7:**
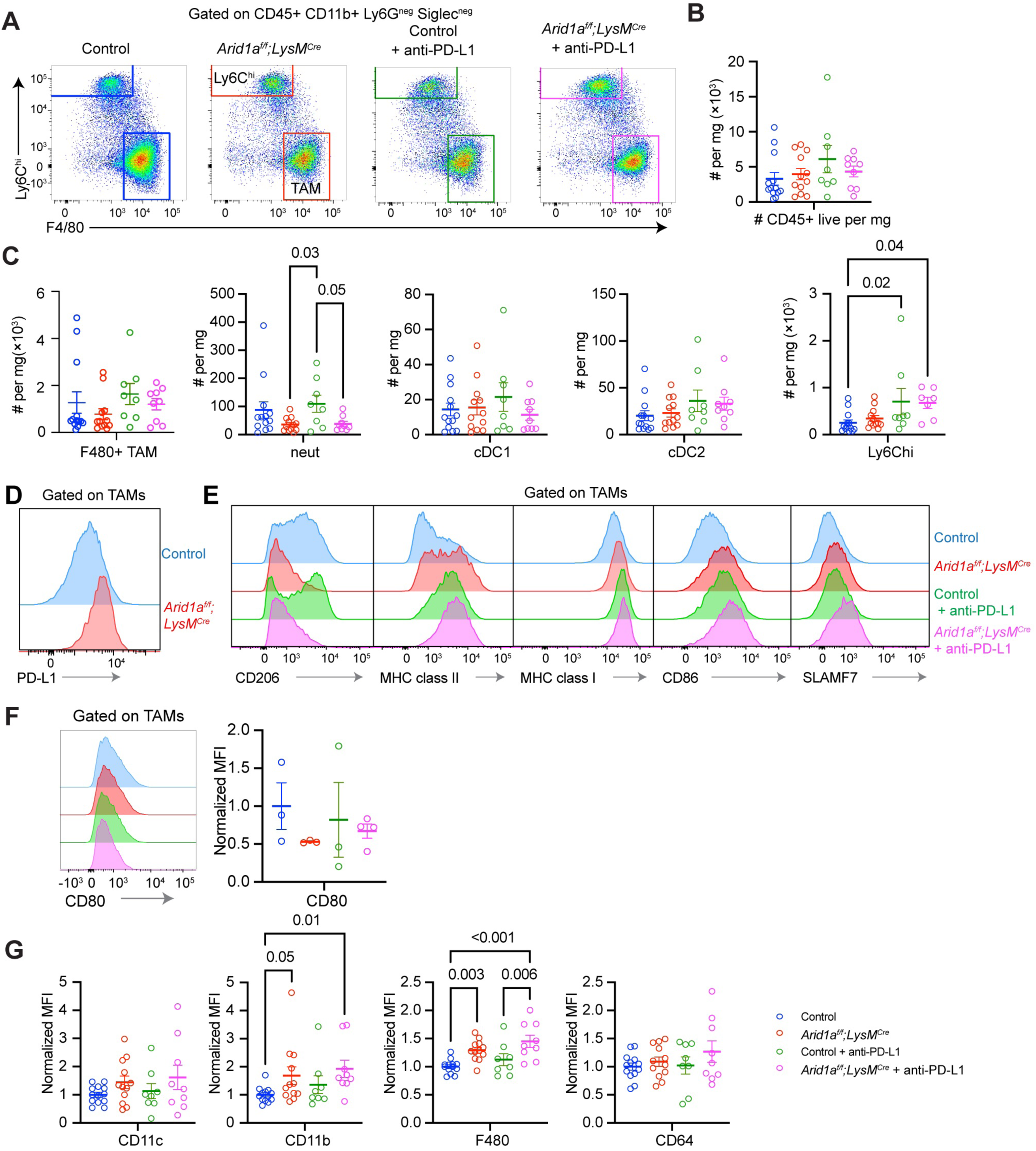
Analysis of tumor-infiltrating myeloid cell phenotype and numbers A. Representative gating of intratumoral TAMs. B. Quantification of the total number of tumor-infiltrating immune cells (CD45+) per mg tumor C. Quantification of the absolute number of the indicated tumor infiltrating myeloid cells. D. Representative histogram for PD-L1 from the indicated groups. E. Representative histogram for the listed cell surface receptors from the indicated groups. F. Quantification of the normalized median fluorescence intensity (MFI) for CD80, accompanied by representative histogram. G. Quantification of normalized MFI of the indicated lineage-markers. Data analyzed by one-way ANOVA between conditions. Shared legend in panel G.

**Supplementary Fig. 8:**
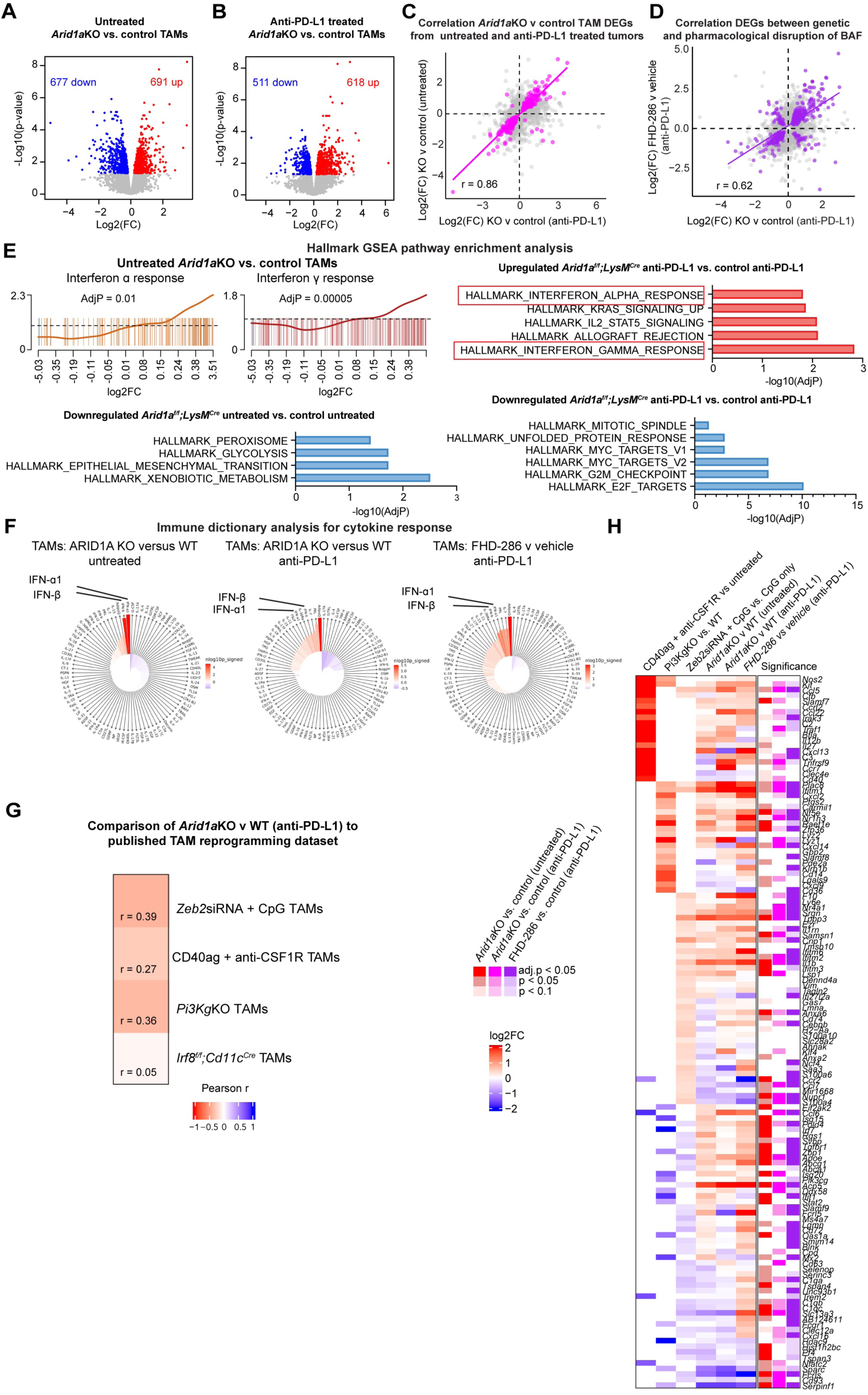
Loss of ARID1A reprograms TAMS transcriptionally A. Volcano plot showing down and upregulated genes in *Arid1a^f/f^;LysM^Cre^* versus control TAMs in untreated groups based on RNA-seq of 6 biological replicates. B. Volcano plot showing down and upregulated genes in *Arid1a^f/f^;LysM^Cre^* versus control TAMs in anti- PD-L1 treated groups based on RNA-seq of 6 biological replicates. C. Correlation plots showing the union of differentially express genes (grey) and highlighting the intersection (magenta) that are differentially expressed genes between *Arid1a^f/f^;LysM^Cre^*versus control TAMs from untreated and *Arid1a^f/f^;LysM^Cre^* versus control TAMs anti-PD-L1 treated tumors. D. Correlation plots showing the union of differentially express genes (grey) and highlighting the intersection (purple) that are differentially expressed genes between *Arid1a^f/f^;LysM^Cre^*versus control TAMs from anti-PD-L1 treated tumors, and TAMs from FHD-286 versus vehicle anti-PD-L1 treated tumors. E. Hallmark pathway analysis of the showing significant up and downregulated pathways in *Arid1a^f/f^;LysM^Cre^* versus control TAM from untreated (isotype-only) or anti-PD-L1 treated tumors. F. Immune dictionary^77^ analysis on differentially expressed genes in *Arid1a^f/f^;LysM^Cre^* versus control TAMs (untreated ie. isotype-only and anti-PD-L1 treated), and FHD-286 versus vehicle (anti-PD- L1 treated). G. Correlation heatmap showing r (Pearson’s correlation coefficient), calculated based on the correlation of log2FC for each gene reported in the TAM reprogramming paper’s supplemental information^22,23,30,94^ (filtered on adjP < 0.05) with the log2FC in the *Arid1a*KO versus control TAM anti-PD-L1 treated dataset. H. Heatmap showing the intersection of genes that were significant (adjP < 0.05) in the reported tables from any of the *in vivo* TAM datasets that were significant in the union of *Arid1a*KO versus control comparisons. Not filtered in direction so it also shows where there is inverse correlation.

**Supplementary Fig. 9:**
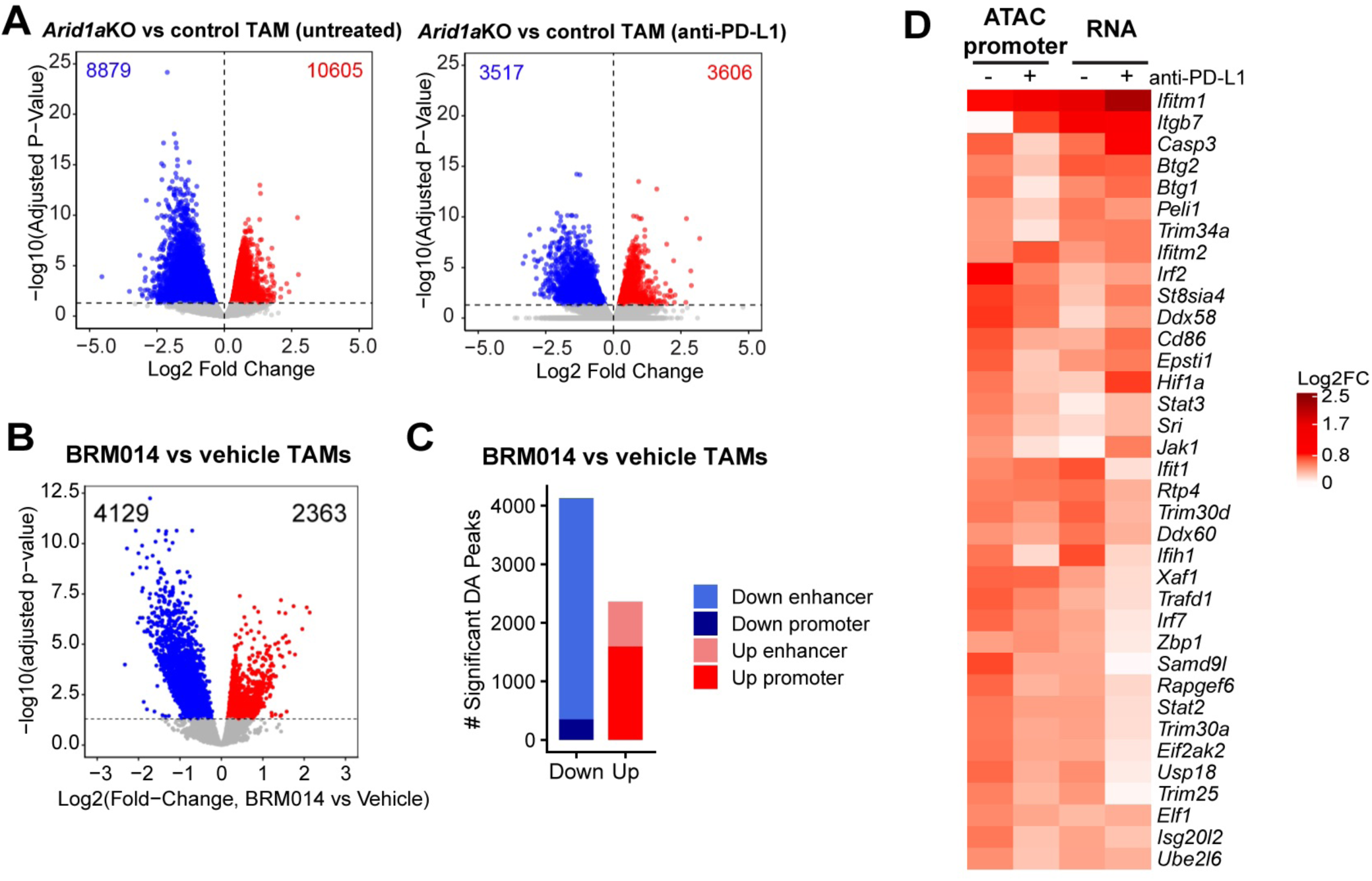
Loss of ARID1A leads to reprogramming of the chromatin accessibility landscape A. Volcano plots highlighting significantly differentially accessible regions between *Arid1a^f/f^;LysM^Cre^* and control TAMs from untreated or anti-PD-L1 treated tumors, as assessed by ATAC-seq. B. Pseudobulk aggregate ATAC-seq analysis of TAMs from scATAC-seq from Multiome assay of MC38 tumors treated with BRM014 + anti-PD-L1 or Vehicle + anti-PD-L1 showing the number of significant differentially accessible sites between conditions. C. Distribution of gene regulatory elements among differentially accessible sites in BRM014 + anti- PD-L1 or Vehicle + anti-PD-L1 from pseudobulk aggregate scATAC-seq Multiome data of TAMs from MC38 tumors. D. Union of ISG promoters that have elevated accessibility and upregulated gene expression (adjP < 0.05) in *Arid1a^f/f^;LysM^Cre^* versus control TAM comparisons.

**Supplementary Fig. 10.**
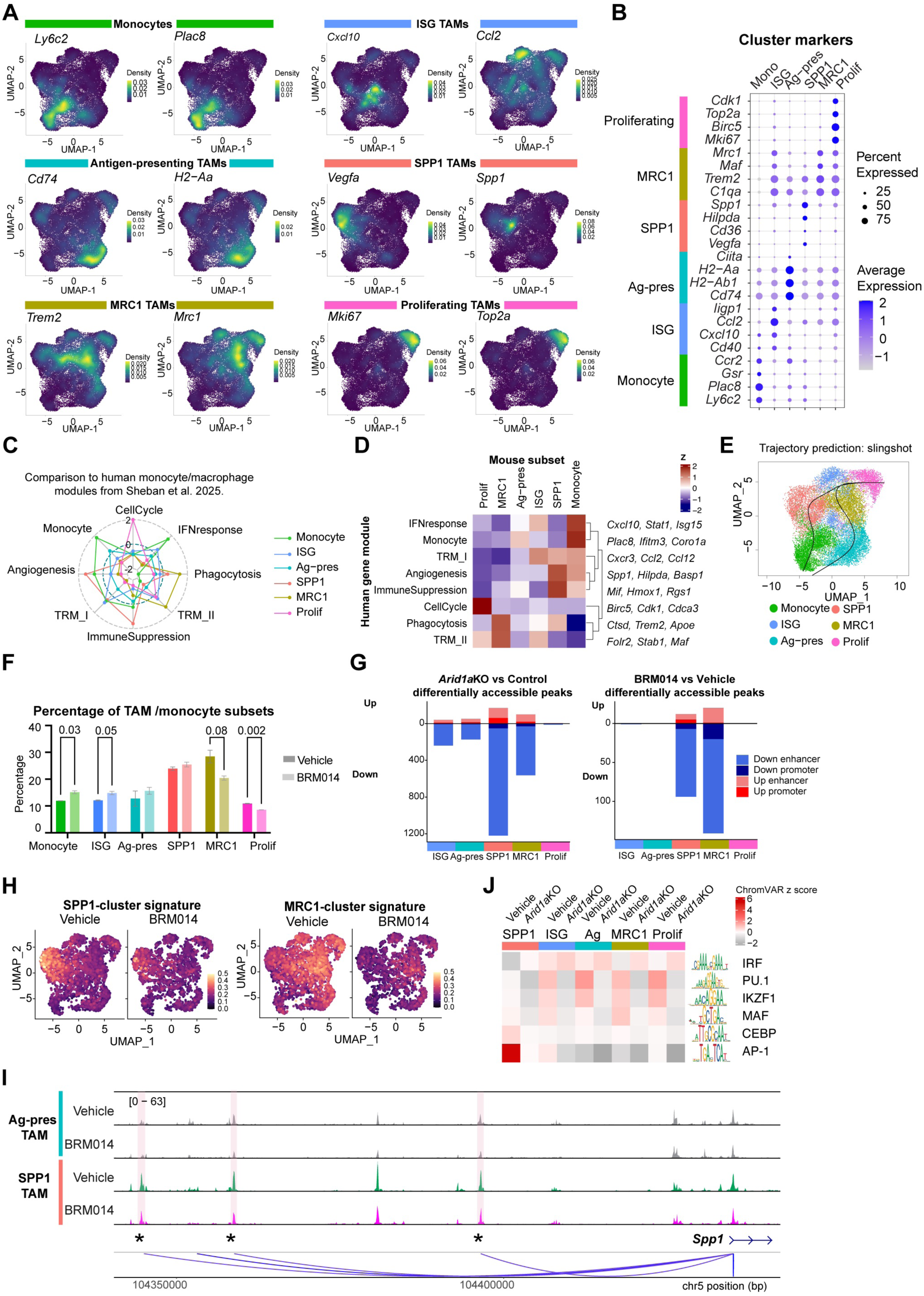
: Multiome and scRNA analysis A. Density plots showing the density of expression of indicated markers across the TAM UMAP in all datasets combined. B. Dot plot showing four markers per cluster that differ in gene expression between TAM substates. C. Radar plot showing z-score of human module signatures from Sheban et al. 2025^30^ within each TAM substate defined in our MC38 data. D. Heatmap showing z-score of human module signatures from Sheban et al. 2025^30^ within each TAM substate defined in our MC38 data. E. Trajectory prediction across TAM subsets based on Slingshot analysis. F. Proportion of cells in each cluster from indicated conditions (all treated with anti-PD-L1) G. Number of significant differentially accessible peaks between BRM014 and vehicle TAMs within each TAM substate showing direction and class (ie. promoter or enhancer), highlighting decreased accessibility in enhancers in MRC1 and SPP1 subsets. H. Expression of signature genes that define SPP1 and MRC1 clusters projected onto the UMAP of the indicated conditions. I. Tracks of *Spp1* enhancers showing differential accessibility between TAM substates and between *Arid1a*KO versus control genotype TAMs. J. ChromVar analysis of motifs between TAM substates and between *Arid1a*KO versus control genotypes.

**Supplementary Fig. 11.**
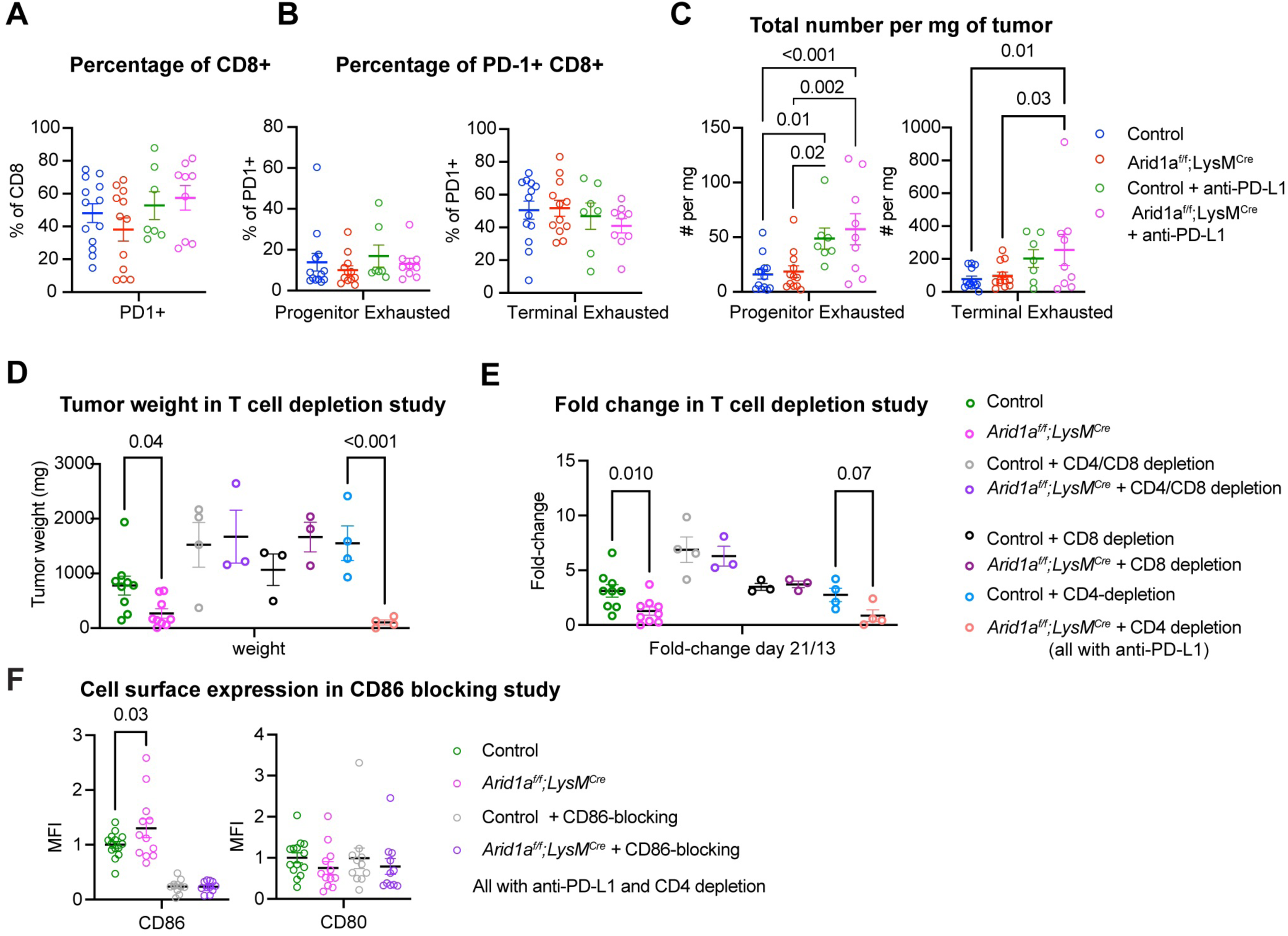
: Loss of myeloid ARID1A leads to changes in intratumoral CD8+ T cells A. Proportion of CD8+ T cells that are PD-1+ of indicated groups (legend as in C). B. Quantification of CD8+PD-1+ T cells expressing exhaustion markers for progenitor (CD8+PD1+SLAMF6+TIM-3^neg^) or terminal exhaustion (CD8+PD1+SLAMF6^neg^TIM-3+) subsets of indicated groups (legend as in C). C. Quantification of the total number of progenitor (CD8+PD1+SLAMF6+TIM-3^neg^) or terminal exhaustion (CD8+PD1+SLAMF6^neg^TIM-3+) subsets cells per mg tumor in indicated groups. D. Weight of anti-PD-L1 treated MC38 tumors of the indicated treatments and treated with CD4, CD8 or a combination of depleting antibodies. E. Fold change from day 21 to day 13 of anti-PD-L1 treated MC38 tumors of the indicated treatments and treated with CD4, CD8 or a combination of depleting antibodies. F. CD86 and CD80 expression on TAMs in CD86-blockade study. Data presented as mean ± SEM, with points indicative of biological replicates. Data analyzed by one-way ANOVA between conditions.

**Supplementary Table 5:** List of antibodies used

| **Antibody target and conjugate** | **Company** | **Catalogue Number** |
| --- | --- | --- |
| Anti-B220-BV785 | BD | 563894 |
| Anti-CD103-BUV737 | BD | 749393 |
| Anti-CD115-BV421 | BioLegend | 135513 |
| Anti-CD11b-PeCy7 | Tonbo Biosciences | 60-0112-U100 |
| Anti-CD11c-BUV737 | BD | 612797 |
| Anti-CD11c-BV650 | BioLegend | 117339 |
| Anti-CD19-BV711 | BD | 563157 |
| Anti-CD19-PE | BD | 557399 |
| Anti-CD206-BV421 | BioLegend | 141717 |
| Anti-CD3-APC-eFluor 780 | Invitrogen | 47-0032-82 |
| Anti-CD4-BUV395 - | BD | 740209 |
| Anti-CD4-BUV737 | BD | 612843 |
| Anti-CD4-PE | BioLegend | 100511 |
| Anti-CD45-BUV737 | BD | 748371 |
| Anti-CD45-BV605 | BioLegend | 100451 |
| Anti-CD45-BV711 | BioLegend | 103147 |
| Anti-CD45-BV785 | Biolegend | 103149 |
| Anti-CD64-APC | BioLegend | 139305 |
| Anti-CD64-PE | BioLegend | 139304 |
| Anti-CD8b-APC | BioLegend | 126614 |
| Anti-CD8a-APC | Tonbo Biosciences | 20-1886-U100 |
| Anti-CD8a-PE | Thermo Scientific | MA517599 |
| Anti-CD8a-BUV395 | BD | 563786 |
| Anti-CD80-APC | BioLegend | 104714 |
| Anti-CD86-BV421 | BioLegend | 105031 |
| Anti-CD86-BV605 | BioLegend | 105037 |
| Anti-CD86-FITC | BioLegend | 105006 |
| Anti-F480-BUV395 | BD | 565614 |
| Anti-F480-BV421 | BD | 565411 |
| Anti-F480-BV785 | BioLegend | 123141 |
| Anti-Ly6C-APC-Cy7 | BioLegend | 128025 |
| Anti-Ly6C-BUV737 | BD | 755201 |
| Anti-Ly6C-BV605 | BioLegend | 128035 |
| Anti-Ly6C-PE | BioLegend | 128007 |
| Anti-Ly6G-APcy7 | BioLegend | 127624 |
| Anti-Ly6G-FITC | BioLegend | 127605 |
| Anti-H-2Kd/H-2Dd -PeCy7 (MHC class I) | BioLegend | 114717 |
| Anti-I-A/I-E BV510 (MHC class II) | BioLegend | 107635 |
| Anti-NK1.1-BV421 | BioLegend | 108732 |
| Anti-NK1.1-PE | BioLegend | 108708 |
| Anti-PD1-APC-Cy7 | BioLegend | 135224 |
| Anti-PD1-BV421 | BioLegend | 135217 |
| Anti-PDL1-BUV737 | BD | 568361 |
| Anti-PDL1-BV650 | BD | 568361 |
| Anti-PDL1-PE | BioLegend | 124308 |
| Anti-Podoplanin-BV421 | BioLegend | 127423 |
| Anti-SiglecF-APC-Cy7 | BD | 565527 |
| Anti-SiglecF-BUV396 | BD | 740280 |
| Anti-SiglecF-BV650 | BD | 740557 |
| Anti-SLAMF6-BV510 (Ly108) | BioLegend | 134606 |
| Anti-SLAMF7-BV785 (CD319/CRACC) | BD | 747993 |
| Anti-Tim3-BV785 | BioLegend | 119725 |
| Anti-Foxp3-FITC | Thermo Scientific | 11-5773-82 |
| Anti-Ki67-BV480 | BD | 566172 |
| Anti-Tox-eFluor 660 | Thermo Scientific | 50-6502-82 |
| Anti-IFNg-PeCy7 | BioLegend | 505825 |
| Anti-GZMB-FITC | BioLegend | 372206 |
| Anti-TNFa-BV711 | BD | 563944 |
| Anti-ARID1A antibody [EPR13501-73] | Abcam | ab182561 |
| Anti-Goat anti-Rabbit IgG (H+L) Highly Cross-Adsorbed Secondary Antibody, Alexa Fluor™ Plus 488 - A32731TR | Invitrogen | A32731 |
| Anti-Goat anti-Rabbit IgG (Heavy chain) Superclonal™ Secondary Antibody, Alexa Fluor™ 647 - A27040 | Invitrogen | A27040 |
| Anti-IFNAR | Leinco | I-401 |
| Anti-mouse IgG1 isotype | Leinco | I-536 |
| Anti-PD-L1 *in vivo* | Bio X Cell | BE101 |
| Anti-rat IgG2b isotype *in vivo* | Bio X Cell | BE0090 |
| Anti-CD4 *in vivo* | Leinco | C1333 |
| Anti-CD8a *in vivo* | Leinco | C380 |
| Anti-rat IgG2b isotype control *in vivo* | Leinco | I-1034 |
| Anti-rat IgG2a isotype control *in vivo* | Leinco | I-1177 |
| Anti-CD86 *in vivo* | Leinco | C2158 |
| Anti-B220-biotin | BioLegend | 103208 |
| Anti-CD11c-biotin | BioLegend | 117304 |
| Anti-CD11b-biotin | BioLegend | 101204 |
| CD4-biotin | BioLegend | 100404 |
| CD49b-biotin | BioLegend | 108904 |
| Anti-TCR$\gamma\delta$-biotin | BioLegend | 118103 |
| Anti-Ter119-biotin | BioLegend | 116204 |
| Anti-CD3 functional | BD | 553057 |
| Anti-CD28 functional | BD | 553294 |
| Anti-AffiniPure Goat Anti-Armenian Hamster IgG (H+L) | Jackson ImmunoResearch | 127-005-099 |
| Anti-CD19-biotin | BioLegend | 115504 |
| anti-CD5-biotin | BioLegend | 100604 |
| Anti-H3K4me3 | Millipore | 05-745R |
| Anti-H3K27ac | Millipore | ABE647 |
| Anti-H3K4me(1) | Abcam | Ab8895 |
| Anti-H3K27me3 | Cell Signaling | C3B11 |

